# Homeotic and non-homeotic patterns in the tetrapod vertebral formula

**DOI:** 10.1101/2024.03.21.586053

**Authors:** Rory T. Cerbus, Ichiro Hiratani, Kyogo Kawaguchi

## Abstract

Vertebrae can be differentiated into five categories along the body axis in tetrapods, with its numerical distribution known as the vertebral formula. The vertebral formula is a principal tool for connecting development and phylogeny [1]. This is largely due to its robust relationship with the conserved clusters of *Hox* genes [2], which exhibit expression boundaries coincident with vertebral divisions [3–11]. One avenue for variations in the vertebral formula is thus through *Hox*-mediated homeotic transformations, which manifest as a relatively fixed sum of adjacent vertebral counts. This expectation is borne out in the mammalian thoracolumbar count [12], but to date, no similar vertebral patterns have been found. Here we conduct a systematic search by generating a large dataset of complete vertebral formulae in a diverse range of tetrapod species and probing the variance of linear combinations of vertebrae. We uncover additional mammalian homeotic patterns, but also unexpected balances between distal vertebrae not comprehensible with *Hox*-mediated regionalization. One distal pattern appears during the progression from theropods to birds, demonstrating its phylogenetic importance. We further show that several vertebral counts correlate with posterior intergenic distances in the *HoxB* gene cluster. By creating a vertebral formula database and mathematically defining patterns, our work establishes a quantitative approach for comparative genomics in morphology.

---

Patterns in vertebrae unite and separate a wide range of animal groups. Based on its location, function, and morphology, an individual vertebra can be classified into one of five different categories: *cervical* (neck, *C*), *thoracic* (rib-bearing dorsal, *T*), *lumbar* (rib-less dorsal, *L*), *sacral* (pelvic, *S* ), and *caudal* (tail, *Ca*). The distribution of the numbers of vertebra in these categories is termed the vertebral formula, which has been studied since the inception of modern comparative anatomy [15–22]. On account of the vertebrae formula alone, a bird can easily be distinguished from a mammal [15], and even orders within mammals are differentiable [1, 12, 23].

Patterns in the vertebral formula are identified in the context of a phylogenetic tree. Most mammals have seven cervical vertebrae while birds exhibit broad variability [17, 18, 21], justifying the significant attention given to the neck. Along with more recent discoveries of patterns further along the body axis, such as the weak invariance of the mammalian thoracolumbar count [1, 12], these patterns have generated important insights and new questions about anatomy [24, 25] and trait evolution [1, 23, 26].

A principal reason for the attention given to vertebral patterns is their manifest but mysterious relationship with the *Hox* genes, a set of strongly conserved homeobox genes that control the body plan of even invertebrates up to Cnidaria [27]. Within tetrapods these *Hox* genes are arranged in a fixed linear order in four clusters [28], and their sequential expression pattern reflects the serialization of the vertebrae, the phenomenon of temporal colinearity [29]. Colinearity is the starting point for the *Hox* code [30], the genotype-phenotype map [31, 32] which serves as a genetic basis for the vertebral formula. While the exact nature of this code and its species-specific manipulation is still a matter of debate [2, 33], the *Hox*-vertebrae pair is a robust system to probe genetic pathways in development.

Following their homeobox namesake, changes in *Hox* gene expression are often associated with shifts of vertebral boundaries, homeotic transformations of one vertebral type into another neighboring type [3–11]. These homeotic shifts, one potential source of variation in the vertebral formula, are thus expected to manifest as anti-correlations and as a relatively fixed sum of counts between adjacent vertebrae. In principle, this yields four conserved sums: *C*+*T, T* +*L, L*+*S*, and *S* + *Ca*. To date, the only well-established example of this is the relatively fixed thoracolumbar count (*T* + *L*) in mammals [12], as well as the possibility that this is behind the deviation from the mammalian ‘rule of seven’ (*C* = 7) for sloths and manatees [26, 34–36] (see also Extended Data Fig. 1). And yet the expectation of *Hox*-mediated homeotic patterns is so strong that it has frequently assumed in the analysis of even extinct species whose genomic sequence is unavailable [1, 37–39]. Here we ask whether vertebral patterns, especially between adjacent vertebrae, are present further along the body axis in not only mammals but other tetrapods as well.

To search for vertebral patterns we construct a broad dataset of complete vertebral formulae for 388 species, representing four tetrapod classes (Mammalia, Aves, Reptilia, Amphibia), 42 orders, and 175 families. This dataset was constructed from a myriad of data sources such as CT scans, photographs of skeletons, scientific literature, and manual inspection of skeletons (see Methods). We here examine the full vertebral count across all tetrapods, in contrast to previous studies of the vertebral formula focusing on subsets of tetrapods, typically mammals, and most often on presacral counts [12, 24, 25, 35, 40, 41]. As represented schematically in Figs. 1a-c, examining these vertebral formulae in the context of a phylogenetic tree yields several types of patterns, both old and new. We observe branches of the tree with nearly constant or strongly varying individual counts, as well as homeotic-type anti-correlations between adjacent vertebrae. We also find an unexpected category, namely correlations between distal regions such as the cervical and sacral, and a balance between the anterior and posterior counts specific to birds. We further probe *Hox*-related vertebral patterns by comparing the variation of the vertebral formula with the *Hox* cluster genetic sequence of the corresponding species, finding regions in the *Hox* cluster for which the intergenic distance correlates with the counts.

**Fig. 1.**
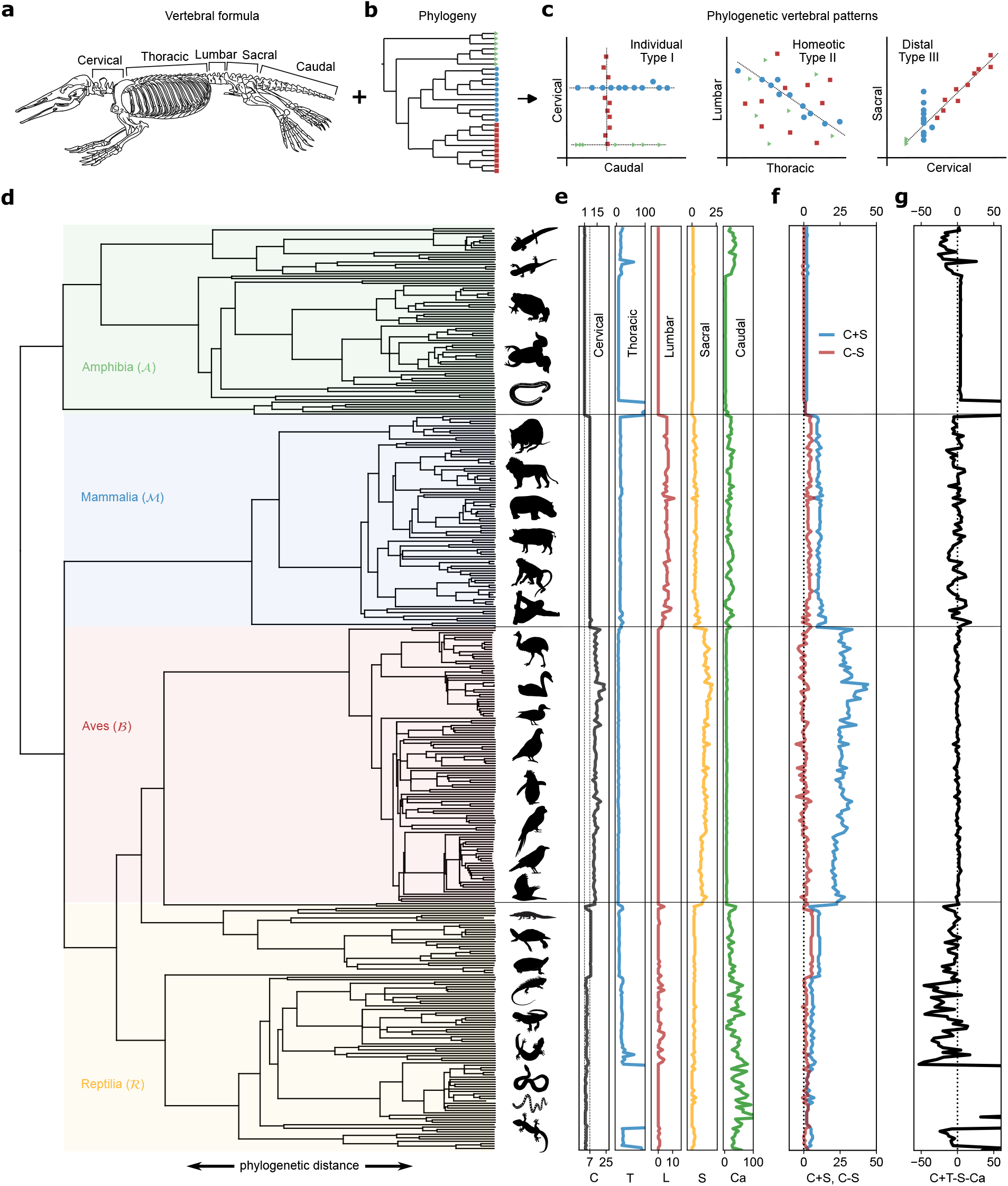
(a)-(c) Schematic of the approach to determine vertebral patterns. (a) Sketch of a platypus (*Ornithorhynchus anatinus*) skeleton reproduced from Cuvier [13]. The numbers of vertebrae from each category yield the vertebral formula. (b) Schematic example of a phylogenetic tree distinguishing three different groups. Analyzing the vertebral formulae in the context of the tree yields the vertebral patterns. (c) We discover three types of constraints. For individual or Type-I constraints, the vertebral count in a single category is nearly constant within a specific branch of the tree. This same category can be very plastic or take on different constant values in other branches. In homeotic or Type-II constraints, adjacent vertebrae are anti-correlated and sum to a nearly constant value. In distal or Type-III constraints, non-adjacent vertebrae are positively correlated and can be numerically balanced. (d)-(g) Plots of the vertebral data arranged according to a phylogenetic tree [14]. (d) A phylogenetic tree containing all the species for which we have the full vertebral count. (We exclude some bird data to avoid unbalancing the tree, but it is used in later analysis.) The tree has been organized so that the four classes of tetrapods, Amphibia, Mammalia, Aves (birds), and Reptiles, are arranged vertically. (e) The individual vertebrae plotted vertically. (f) Vertical plots of two novel constraints and plasticities discovered with our approach. *C* − *S* is a global constraint found when comparing all tetrapods. *C* + *S* is a plasticity found within birds. (g) Vertical plot of a novel constraint, *C* + *T* − *S* − *Ca* found for all birds.

## Patterns in the vertebral formula

There is a variety of nomenclature to describe patterns such as those we treat here, with many terms implying a particular origin to the pattern [42]. Here we do not attempt to determine a pattern’s origin, such as whether it is due to a shared developmental program, physical restriction, stabilizing selection, or phylogenetic inertia. With the null hypothesis that vertebral counts are free to vary and independent, we label any significant pattern with low variation a “constraint”, and any significant pattern with large variation a “plasticity” [43].

We collect data into a two-dimensional array *V*_*i j*_ with five columns corresponding to the five vertebral categories. Each row corresponds to a different species, so that the vertebral formula for a species *i* is given by (*C*_*i*_, *T*_*i*_, *L*_*i*_, *S* _*i*_, *Ca*_*i*_). To search for patterns in the tetrapod phylogenetic tree, we query TimeTree [14], a consensus tree, with the list of species included in our dataset and produce the tree in Fig. 1d. We plot the individual components *V*_*i j*_ for each species *i* to the right of this tree (Fig. 1e). In addition to known patterns such as *C* ≃ 7 in Mammalia [12] and *C* = 1 in Amphibia (the atlas) [19], we also observe a high variance of *Ca* in Mammalia, Reptilia, and some orders of Amphibia. On the other hand, patterns involving multiple vertebrae such as the combined thoracolumbar (dorsal) vertebrae in mammals, *T* + *L*, as well as the novel constraints that we we describe in detail later (Figs. 1f,g) are easy to overlook in plots of single vertebrae.

To mathematically identify constraints and plasticities, we first iterate through all branches *B* with at least *N* = 20 tips and extract the subset of *V*_*ij*_ corresponding to this branch, 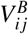. We then calculate the variance of linear combinations of categories,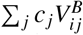, where *c*_*j*_ are the coefficients, to determine if these qualify as constraints or plasticities. In this notation, if we restrict attention to the Mammalian branch *B* = M, then the coefficients for considering only *C* will be *c*_*j*_ = (1, 0, 0, 0, 0), and the average (⟨⟩) of the cervical count in Mammalia will be given by 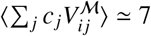.

We identify patterns by comparing the variance of 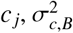, with the total variance of that branch, 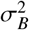, and the variance of the same *c*_*j*_ for all species outside that branch, 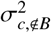 . We determine *c*_*j*_ which are constraint or plasticity candidates by utilizing principle component analysis (PCA) [47], which rotates the original 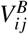 and sorts the new dimensions by variance. We identify the earlier dimensions representing higher variance as plasticities, and the later dimensions, representing lower variance as constraints, by setting a threshold criteria. For a particular PCA component *c* _*j*_, if 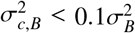 or 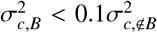, we call *c*_*j*_ a constraint, andif 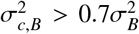 or 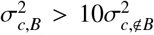, then we call *c*_*j*_ aplasticity. We also search manually for constraints and plasticities by computing 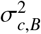 for all linear combina-tions *c*_*j*_ with integer coefficients between [-3,3] and combine these results with those from our PCA analysis if they also satisfy the same threshold criteria (see Methods). We then screen them with a standard phylogenetic test to ensure they are not simply products of the tree structure (PIC [45], see Methods), and determine the earliest branch possessing this constraint or plasticity. An annotated tree with all patterns detected in this way is shown in Extended Data Fig. 2a, and a complete table of discovered patterns is shown in Extended Data Fig. 4.

Our method identifies well-known patterns in the vertebral formula (Fig. 2a) as well as additional plasticities and constraints, the latter of which come in three flavors (Figs. 1c, 2b). The first (I) and most familiar is when a single vertebra is nearly fixed, as with the highly conserved mammalian cervical count (*C* ≃ 7, Fig. 2c). As Figs. 2c,d show, a vertebra can be a constraint in one branch and plastic in another; we find plasticities in a single vertebra such as the highly variable caudal count in mammals, reptiles, and Urodela (salamanders) (Fig. 2c).

**Fig. 2.**
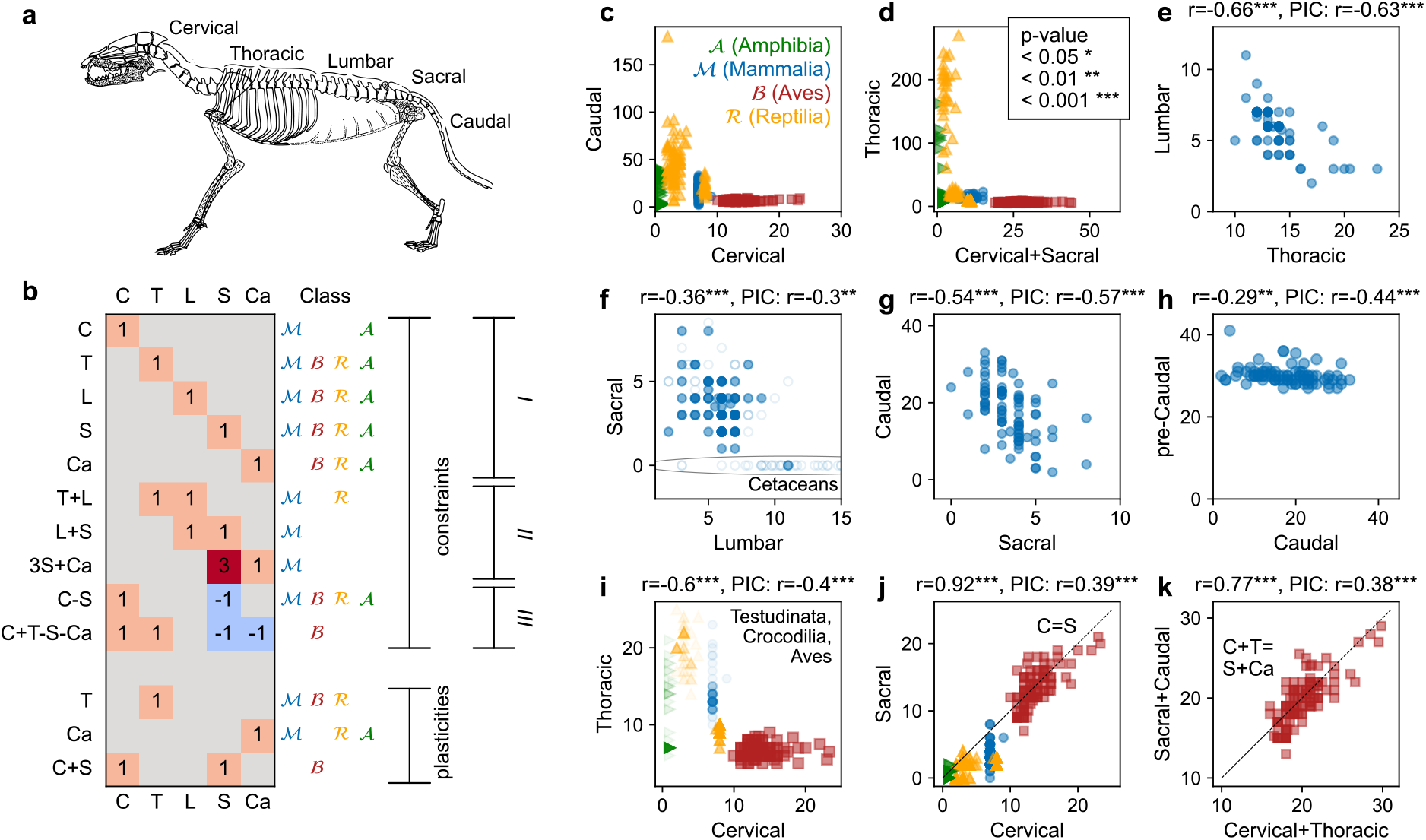
Representative constraints and plasticities. (a) Sketch of a prototypical Mammal redrawn from Owen [44]. (b) Table with representative constraints and plasticities, including their formula, the tetrapod class(es) within which they are found, and the constraint category (single: I, homeotic: II, distal balance: III). Several novel constraints (plasticities) have been discovered. (c)-(k) Plots of representative constraints and plasticities (re-)discovered using our method. For combination constraints (II, III) we calculate not only the Pearson correlation coefficient (*r*) between vertebral categories but also between their Phylogenetic Independent Contrasts (PIC) [45]. In (c) and (d) we plot vertebral counts which can be constraints for some Classes and plasticities for others, yielding an L-shaped plot. (e)-(g) Mammalian homeotic patterns (II) including the previously known *T* + *L* and the newly discovered *L* + *S* and 3*S* + *Ca*. In (f) we also plot genus-averaged data from partial vertebral data [25] and the full vertebral data of Cetaceans [46] as open circles with higher transparency (up to *L* = 15). Two additional homeotic patterns (II) are found for Testudinata (h) and Testudinata with Archosauria (i). In the background of (i) we additionally plot all tetrapods with a higher transparency. Excluding Amphibia and snakes yields significant correlation between *C* and *T* (*r* ≃ −0.84^***^). (j)-(k) Plots of two prominent distal balance constraints (III). (j) The *C* − *S* constraint spans all tetrapods, while in (k) the *C* + *T* − *S* − *Ca* is unique to birds.

A second (II) category of constraints is the anticorrelated adjacent vertebrae, a likely manifestation of a homeotic transformation [1] as the addition of two adjacent vertebral counts is conserved. The only previously known example in this category is the mammalian *T* + *L* [12] (Fig. 2e). We have found additional weak anti-correlations within Mammalia further along the body axis corresponding to constraints on *L* + *S* (Fig. 2f), and a stronger constraint corresponding to 3*S* + *Ca* (Fig. 2g). We note that a homeotic transition between the lumbar and sacral vertebra has been discussed in the context of intra-species variability related to different modes of motion [24, 25]. The peculiar coefficient on the constraint 3*S* + *Ca* (Fig. 2g) suggests that a single sacral vertebra can be substituted by three caudal vertebrae. An unequal conversion by homeotic transformation may be taking place to increase the variability in the caudal vertebrae in mammals. These results suggest that the entire mammalian axis obeys the homeotic transition rule, consistent with the *Hox*-vertebrae code. We observe a near-constant pre-caudal count compared with the caudal count in mammals (Fig. 2h), but with a small yet significant negative correlation between them. Outside Mammalia there are other examples of category II constraints such as the *C* + *T* in Testudinata and Archosauria (birds and alligators, Fig. 2i), which we also detect when including all reptiles and mammals, especially those not obeying the *C* = 7 rule (sloths and manatees, Extended Data Fig. 1).

In stark contrast to these two constraint categories, our method uncovers a novel third category (III) which is a balance between distal (non-adjacent) vertebrae, yielding a positive correlation as seen in Figs. 2j,k and Extended Data Fig. 4. Such correlations between distal regions are intriguing since they are not homeotic transformations and cannot be reconciled with the current understanding of the *Hox* code [1]. We next discuss two prominent examples from this constraint category.

### Correlations between distal vertebrae

At the phylogenetic scale of the tree, we discover that *C* − *S* is a constraint, with *C* ≃ *S* over a wide range of *C* and *S* . In Fig. 1f we plot *C* − *S* and *C* + *S* with the vertical location corresponding to the tree in Fig. 1d. As Fig. 2j also shows, while within some classes there are significant departures from *C* ≃ *S*, such as in Testudinata (*C* ≃ 8) and Mammalia (*C* ≃ 7), especially in tree sloths [34] (see Extended Data Fig. 1), at the tetrapod phylogenetic scale we find that small *S* follows small *C* (Amphibia) and large *S* follows large *C* (Aves). Within Aves and Reptilia, we find that *C* ≃ *S* holds even when *C* and *S* can individually vary substantially. Indeed we also find that the combination *C* + *S* is plastic within Aves (Fig. 2d), highlighting the agreement between *C* and *S* therein.

Another exceptional constraint identified by our method is a balance in the anterior (*C* + *T* ) and posterior (*S* +*Ca*) vertebral counts of birds. It is well known that for birds *C* can vary considerably from less than 10 to more than 23 [18, 41]. And yet it is manifest from Figs. 1g, 2j that *C* is strongly correlated with *S* in Aves, and we also find that *T* and *Ca* vary much less and are weakly correlated. When combined, the anterior *C* + *T* and posterior *S* + *Ca* have a significant correlation (*r* ≃ 0.779, p-value ≃ 4.60 × 10^−41^). Moreover, *C* + *T* − (*S* + *Ca*) ≃ 0, indicating an anterior-posterior balance (Figs. 1g, 2k).

We further analyze this bird constraint by including additional data omitted in Figs. 1,2. We analyze a total of 178 unique bird species (196 total data entries) representing 29 unique orders. In addition to a large portion of the Aves data coming from a book by Eyton in 1875 [22], we examined 81 bird skeletons at the Abiko Bird Museum in Abiko, Japan, to accurately count especially *S* . All data adhere to the bird constraint *C* + *T* ≃ *S* + *Ca* regardless of the source (Extended Data Fig. 3). We show several of these skeletons, and a historic sketch, in Figs. 3a-f. In Fig. 3g we show the vertebral formulae for birds from 21 different orders, as well as examples from extinct theropods and other tetrapod classes. The birds, the bat (*Myzopoda aurita*), and *Archaeopteryx* show a conspicuous anterior-posterior balance. We find that the departure from the bird constraint for each order, quantified as 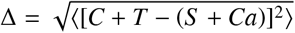, is comparable to the departure averaged over all orders, ⟨Δ⟩ ≃ 1.80, with the exception of Casuariiformes (Δ ≃ 5.70), an order with only two data from Emu (see Extended Data Fig. 3a).

**Fig. 3.**
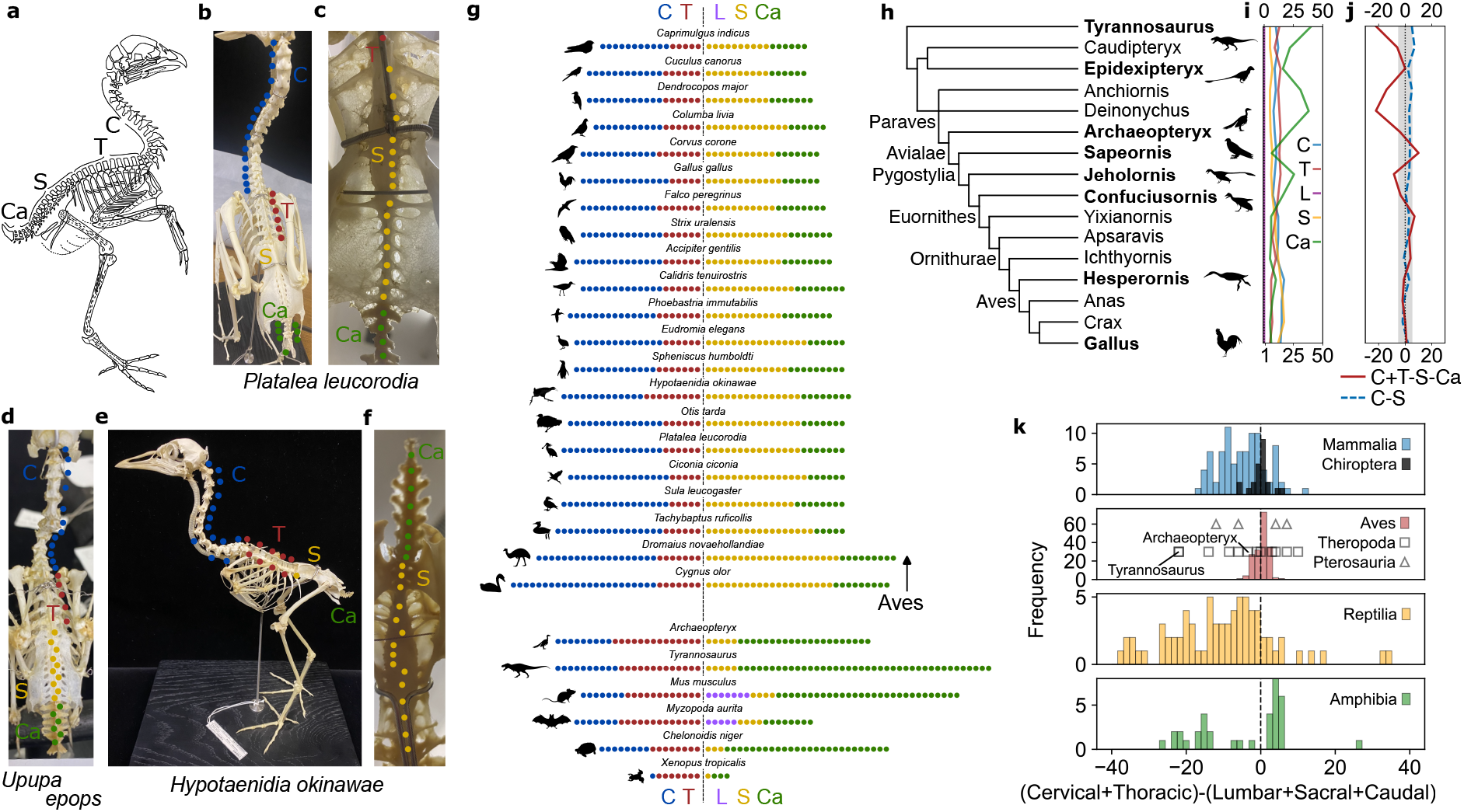
Analysis of the bird constraint. (a) Sketch of a prototypical bird redrawn from Owen [44]. (b)-(f) Images of the skeletons of *Platalea leucorodia* (Eurasian spoonbill), *Upupa epops* (Eurasian hoopoe), and *Hypotaenidia okinawae* (basionym: *Rallus okinawae*, Okinawa rail) from Abiko bird museum indicating *C, T, S*, and *Ca*. (g) Plot of the vertebral formula represented by a number of different colored circles. Here we show birds from 21 different orders in Aves, as well as extinct Theropod species and extant species from the three other tetrapod classes. The vertical dashed line is an anterior-posterior boundary located after the thoracic (*T* ). Birds show a balance between the two sides. (h) Cladogram of extinct and extant theropods extracted from a strict consensus tree [48]. (i) Plot of the individual vertebral counts corresponding to the tree. (j) Plots of *C* − *S* (dashed line) and *C* + *T* − (*S* + *Ca*) for the species on the left. The gray band denotes three standard deviations of *C* + *T* − (*S* + *Ca*) around the average value of ≃ 0.084 for extant birds. All taxa conform to the tetrapodian *C* − *S* constraint while more phylogenetically distant theropods do not conform to the bird constraint. (k) Histograms of (*C* + *T* ) − (*L* + *S* + *Ca*) differentiated by tetrapod class to test a “modified” bird constraint (*L* = 0 for Aves). All other groups of tetrapods save the putative flyers such as birds, bats (Chiroptera), and Pterosauria deviate strongly from the modified constraint.

With the expectation that this bird constraint may also shed light on the evolution of birds, we also investigate the vertebral formula in the clade Theropoda, which includes extinct non-avian dinosaurs such as *Tyrannosaurus rex*. In Fig. 3h we show a cladogram extracted from a strict consensus tree [48]. In Figs. 3i,j we plot the individual vertebral counts, the tetrapod constraint *C* − *S* (dashed line), and the bird constraint *C* + *T* − (*S* + *Ca*). We find that the tetrapodian *C* − *S* constraint is even satisfied by the dinosaur *Tyrannosaurus rex*. The deviation from the bird constraint, on the other hand, increases as the phylogenetic distance from modern birds increases, primarily due to the caudal count [9]. One exception to the trend is *Epidexipteryx*, a taxon which is notably difficult to place phylogenetically [48, 49]. Likewise, the deviation for *Sapeornis, Jeholornis*, and *Confuciusornis* is not monotonic in phylogenetic distance according to this tree [48]. The phylogenetic placement of these three taxa is often shuffled in other cladistic studies [9, 48, 50–54], indicating that the bird constraint may help in resolving these difficulties.

To seek the possible meaning of the bird constraint, we modify our comparison of birds with other tetrapod classes by including *L*, which is always 0 in Aves. Developmental studies suggest that the synsacrum of birds may contain somites that originally had affinity with lumbar vertebrae [3], so in Fig. 3k we plot histograms of a modified bird constraint: (*C* + *T* ) − (*L* + *S* + *Ca*). When we test another extant group of flyers, Chiroptera (bats) [55, 56], and an extinct group of putative flyers, Pterosauria [53, 57– 61], we find that the mea n deviation from the modified bird constraint, 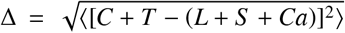, is ≃ 3.27 and ≃ 7.83, respectively, while it is ≃ 8.29 in mammals as a whole. Interestingly, the theropod species that are considered flightless typically show a large deviation, whereas in likely flyers or gliders the deviation is small. We conjecture that the bird constraint is a result of selection for flight biomechanics, which will be important in the discussions of the flight capabilities of extinct theropods and pterosaurs.

### Genetic patterns

We next ask whether any of the vertebral variations, in particular the homeotic patterns (category II), can be directly connected with the *Hox*-vertebral code using a similar many-species comparison [31, 32]. In Figs. 4a,b we review the state of the art regarding the coincidence between *Hox* gene regionalization and vertebral boundaries. We combine the results of a literature survey of species-specific studies on the correlation between individual *Hox* gene’s anterior expression boundaries or perturbations (knock-ins and knock-outs) and homeotic vertebral transitions [3–5, 7–11]. Fig. 4b visually demonstrates the overall colinearity between the *Hox* genes and the vertebrae. As the somites destined to become vertebrae are formed from head to tail, the *Hox* genes are expressed in parallel and in all clusters from paralogous gene group (*PG*) 1 to 13. The colinearity demonstrated in Fig. 4b suggests which genetic region to probe to understand the homeotic vertebral constraints (category II) in particular since these are likely due to homeotic transformations. However, the colinearity is not perfect, and some genes such as *PG9*, correlate with many vertebral transitions. Moreover, Fig. 4b does not portray magnitude, and there is evidence that posterior genes control anterior ones, the phenomena of “posterior prevalence” [29].

**Fig. 4.**
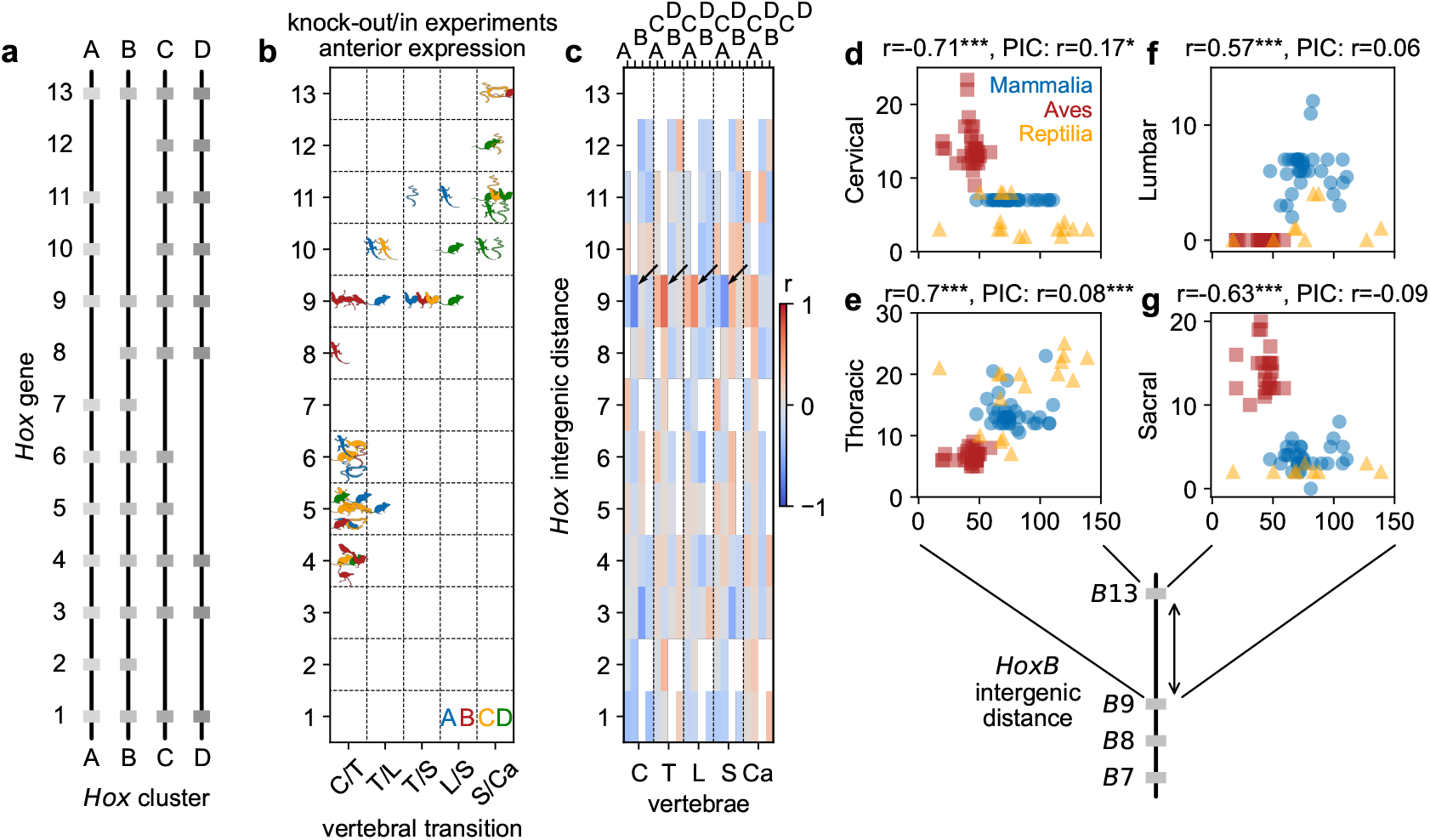
Plots of the correspondence between the *Hox* genes and vertebrae. (a) Schematic inventory of the *Hox* genes in the four clusters present in the putative tetrapod ancestor [28]. (b) Summary of knock-out/in experiments and experimentally determined anterior expression boundaries of individual *Hox* genes that correlate with vertebral transitions [3–5, 7–11]. The species used in the study is indicated by the data point shape, and the specific *Hox* cluster is denoted by the color. Studies often focus on specific *Hox* regions, and so, for example, the higher density of data points near the anterior *Hox* genes does not represent a stronger correlation. A general colinear trend is observable wherein anterior and posterior *Hox* genes correlate with anterior and posterior vertebrae respectively. Also consistent with the data is a “posterior prevalence” [33] in which posterior genes, such as *HoxPG9* (paralogous group), also correlate with anterior vertebrae. (b) A plot of the Pearson correlation coefficient (*r*) between the *Hox* intergenic distances and the vertebral numbers for all classes. The two strongest correlations we find are between the *HoxB9-B13* intergenic distance and the *C* and *T* counts, which we indicate by arrows. (d)-(g) *C, T, L*, and *S* vs. intergenic distance between *HoxB9* and *HoxB13*. The relationship between *B9* and *C* and *T* is anticipated by the work of Moreau et al. on Avian forelimbs [11]. Because *HoxB13* is missing in many Amphibians they are noticeably absent from (c)-(d) [28].

The apparent species-independence, while a testament to the ubiquity of the *Hox*-code [29], also begs the question as to how it can mediate transformations in a species-specific way. While the *Hox* genes themselves are conserved, the sequence around them is highly variable in terms of transposable elements and intergenic distance [10]. Recent work has shown that the number of CCCTC-binding factor (CTCF) sites between *HoxD* genes can significantly modify their expression in mouse gastruloids [62], suggesting CTCF binding sites as a prime candidate for the regulation of the vertebral count through the *Hox* genes. The CTCF binding sites can serve as anchors for loops in chromatin, which may modify the timing and amount of transcription [62, 63].

Experimental information about CTCF binding sites is mostly restricted to mammals and a few model organisms, thus prohibiting us from directly testing this with diverse tetrapod species. Instead, we use the intergenic distance as a proxy for the number of CTCF binding sites. We justify this approach by comparing the size of *Hox* clusters in several model organisms with the number of experimentally determined CTCF binding sites therein and found a significant correlation (*r* ≃ 0.81, p-value ≃ 1.53 × 10^−4^, see Extended Data Fig. 7a).

Using genome assemblies available from NCBI, we generate a database of species for which we have at least partial vertebral formulae and for which the *Hox* clusters are each located on a single chromosome or scaffold. We determine the *Hox* gene locations either from existing annotations or by performing a *BLAST* search [64], yielding a combined database of *Hox* intergenic distances and (at least partial) vertebral formulae with 107 species across the four tetrapod classes. We have not included snakes in this analysis since other work has shown that their unusual vertebral count is largely controlled by other factors [65].

We compare the intergenic distances with the individual vertebral counts for all species and produce the Pearson correlation matrix shown in Fig. 4c (we show the correlation within classes in Extended Data Fig. 6). Within our dataset, the strongest and most significant correlations are between the *HoxB9-13* intergenic distance and the *C, T, L* and *S* counts (Figs. 4d-g). The signs of the correlations are opposite, in accord with the anti-correlation between *C*/*T* and *L*/*S* suggestive of a homeotic transformation shown in Figs. 2f,i. Even though mammals have an essentially fixed *C* count, they still fall along the global trend, much as they did with the correlation between *C* and *S* (see Fig. 2d and Extended Data Fig. 1). The correlation between *C, T*, and *HoxB9* is consistent with the previous finding that the forelimb, typically located at the *C*/*T* transition, coincides with the position of switching in the expression from *HoxB4* to *HoxB9* in several bird species [11].

In general, we did not find correlations that support the idea that the vertebral formula is controlled by the *Hox*-cluster genetic structure. Within the correlations between the intergenetic distance and the vertebral counts, only a few survived the phylogenetic test (PIC). However, just as there are branch-specific vertebral patterns (Fig. 2), we found scattered correlations specific to certain tetrapod classes (Extended Data Figs. 6,7). In fact, the most significant correlations between vertebrae and the intergenic distances, for *HoxB9-B13*, seem to align with the picture of posterior prevalence [33]. Our findings suggest a need to look beyond only co-location to also probe the respective weights of the *Hox*-vertebral correlations.

## Discussion and conclusion

By generating a database of complete vertebral formulae for tetrapods, we identified three main patterns; constant single count (I), homeotic (II), and distal correlations (III). We found that the vertebral formula is largely constructed according to a homeotic pattern within mammals, but largely not in other tetrapod classes, casting doubt on a universal *Hox*-vertebral code. Moreover, we identified correlations between distal vertebrae for tetrapods and birds that are not comprehensible within the current understanding of the *Hox*-vertebral relationship.

Important next steps will be to investigate the origin or nature of the vertebral patterns. Direct investigation through developmental perturbations can ascertain whether constraints are developmental or evolutionary [42]. For example, manipulating the expression of *Hoxb9* in chickens, which may affect the cervical count [11], could permit a test of the *C*/*S* distal constraint. Alternatively, patterning in organoid models could provide a more accessible system for manipulating *Hox* gene expression to test its relationship with vertebral patterns [62, 66].

Indirect approaches, such as correlating cooccurring features, may offer additional insights. Examples include the apparently coincidental (pleiotropic) relationship between the cervical count and mammalian cancer [67] or mammalian diaphragm [68], or the consistent vertebral columns in *Anura* despite significant phylogenetic diversity (see Fig. 1). The possible connection between avian flight and the unique bird constraint exemplifies this approach. It will be also important to consider whether patterns arise at the point of segmentation or much later, as may be the case for the avian synsacrum [69]. Furthermore, exploring intraspecies vertebral count variability can yield valuable insights. Investigating intra-species variations of lumbosacral counts in mammals has yielded potential relationships with mobility and environment [24, 25]. Likewise, within the mammals that disobey the ‘rule of seven’ or which have entirely lost their sacral vertebrae, the intra-species variations suggest a homeotic origin [34, 35, 46] (see also Extended Data Fig. 1). We anticipate that probing intra-species variations in birds relative to their unique constraint may also shed light on their origins. Finally, extending our analysis to encompass the entire vertebral tree including fish [70] should enrich our phylogenetic perspective on vertebral patterning.

## Data availability

The tables of vertebral formulae, *Hox* intergenic distances, and other relevant information will be deposited in an online repository.

## Code availability

Code for the implementation of PIC and analysis of vertebral and intergenic data is available along with the data in the online data repository [71].

## Acknowledgments

We thank Y. Odaya and the Abiko City Museum of Birds for their hospitality and help in viewing their bird skeletons, and Shigeru Kuratani for critically reading the manuscript. We also thank W.M. Cerbus and E.D. Cerbus for their help in redrawing the sketches from Cuvier and Owen [13, 44]. This work was supported by JSPS KAKENHI Grant Numbers JP19H05795, JP19H05275, JP21H01007, and JP23H00095 (to K.K.).

## Author contributions

R.T.C. and K.K. designed and implemented the study. R.T.C. collected and analyzed the data. R.T.C., K.K., and I.H. wrote and edited the manuscript.

## Methods

### Vertebral categorization

In order to compare the vertebral formula between different classes of tetrapods, it was necessary to identify categories of homologous vertebrae. Different nomenclatures and conventions exist for different classes and even orders, and so we determined a consensus definition for each vertebral category. The cervical (*C*) bones were defined as those vertebrae between the skull and the first rib-bearing vertebrae, including the axis and atlas. In mammals and amphibians, this definition leaves no ambiguity as to the identity of the cervical vertebrae. In some reptiles (e.g., Crocodilia) and birds, however, other authors have identified the cervical vertebrae as all vertebrae between the skull and the forelimb. We next define the thoracic vertebrae (*T* ) as all rib-bearing vertebrae in the dorsal region, anterior to the pelvis or sacrum. This includes free or “false” ribs not attached to the sternum and excludes vertebrae with ribs that are fused with the synsacrum in birds [22]. The lumbar (*L*) vertebrae are all remaining dorsal vertebrae posterior to the rib-bearing thoracic and anterior to the pelvis or sacrum. With this definition birds and amphibians and many reptiles do not have lumbar vertebrae but most mammals do. We define the sacral vertebrae (*S* ) as those vertebrae which are (typically) fused together to form the pelvic region to which the hindlimbs are attached. Some authors have made further categorizations of vertebrae within the fused synsacrum of birds [3], but we go no further than to identify them as sacral. In this, we essentially follow Eyton’s identification [22]. Identifying the individual sacral vertebrae in birds is especially challenging due to their reduced size and complex ossification (see Fig. 3 and Extended Data Fig. 8). For snakes, which do not have a hindlimb, the sacrum is identified with the cloacal when possible [65]. Finally, the caudal vertebrae (*Ca*) are identified as all free vertebrae posterior to the sacrum. We counted the bird’s fused pygostyle as a single vertebra, and following Ro č kov á et al. [72] we assigned a caudal count of 3 for Anura.

### Extant data collection

The vertebral counts for extant species were obtained from a variety of sources, including literature where the counts are already given or could be surmised from figures, CT scans, and images of skeletons from online repositories (Morphosource, Digimorph, Sketchfab), and from manual inspection of bird skeletons at the Abiko Bird Museum in Abiko, Japan. A full list of sources can be found in the electronic Supplementary Material along with the vertebral data (Supplementary Data #?). The precise age and sex of the majority of the specimens are unknown, but based on overall size we presume that most of the specimens are adults. Some of the counts were taken from studies of embryos, for which the authors identified which vertebral type the somites eventually became. We also provide several example images of skeletons, X-rays, and CT data in Extended Data Figs. 8, 9.

### Algorithm for finding vertebral patterns

We search for vertebral patterns within the vertebral data *V*_*ij*_ in the context of an existing phylogenetic tree [14]. For each branch *B* of the tree with at least *N* = 20 tips (species), we identify which linear combinations *c*_*j*_ of the vertebral data columns have large or small variation 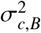 compared to both the total variation of this branch 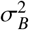 and the variation of the same linear combination for all species (tips) outside this branch 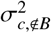. Because the number of linear combinations *c*_*j*_ to test is in principle infinite, we use two tools to focus our search. First, we use principle component analysis (PCA) [47] to reorganize and rotate the vertebral data and sort it by variance. PCA will naturally pick out features (*c* _*j*_) with large and small variance, making it ideal for this task. Second, we test all linear combinations with integer coefficients ranging from -3 to 3. This yields a total number of 7^5^ = 16807 possible *c*_*j*_, which reduces to 8403 unique *c*_*j*_. We compile a list of *c*_*j*_ from both techniques and for each branch *B* that satisfy the criteria for either a constraint, 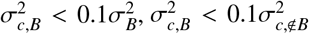, or a plasticity 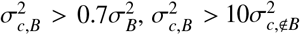. We provide this master list in the Supplementary Material for the criteria we use. For this *c*_*j*_ where possible (a constraint with more than one non-zero coefficient), we also tested whether the Pearson correlation coefficient *r* determined after re-normalizing the counts using the method of phylogenetic independent contrasts (PIC, see Sec. 8) [45], is greater than some threshold value (here *r*_PIC_ > 0.3). We combine the results from the integer *c* _*j*_ and the *c* _*j*_ determined by PCA, which are typically not integers, by rounding the PCA *c* _*j*_ coefficients to the nearest integer. For the combined list, we then proceeded to identify all unique constraints and plasticities and determine the highest (earliest) node on the tree at which they hold. Some constraints appeared independently in the tree, such as the constraint on *T* + *L* in both Mammalia and Testudinata. This final list of constraints and plasticities are labeled in Fig. 1a, tabulated in Extended Data Fig. 4, and plotted in part in Fig. 2. In Extended Data Fig. 5a we show the total number of constraints and plasticities identified as a function of these different variance thresholds. We provide the Python code used to analyze the vertebral data in a repository [71].

### Phylogenetic Independent Contrasts

It is possible that correlations between the phenotypic characters of different species may be manifestations of the structure of the phylogenetic tree to which the species belong [45]. The placement of the species in the tree is itself determined by morphological and genetic differences. For the correlation of the characters in question to be significant, their relationship must differ from what is expected purely by their respective phylogenetic distance. Put another way, we could imagine the two characters under scrutiny take some original value at the root of the phylogenetic tree, and we let the value of these characters undergo a random walk along the branches. If their final values at the tips are correlated in the same way as we observe in the same measure as the actual data, our correlation is not significant.

We use the method of phylogenetic independent contrasts (PIC) originally proposed in [45] to deal with this potential problem. In this method, the character values at the tree tips are subtracted from their nearest neighbor and normalized by their respective phylogenetic distance. The contrasted tips are then merged using a weighted average and the process is repeated until all tips are accounted for. A tree with *N* tips will thus yield *N* − 1 contrasts. The subtraction or contrast is designed to remove the shared influence of the tree before their common branching point, and the normalization makes it easier to compare contrasts with different (original) branch lengths. We used a relatively low threshold for the Pearson correlation coefficient of *r* = 0.3 after the PIC for screening the vertebral patterns belonging to categories *II* and *III* (it is not possible to test category *I* constraints with PIC). We wrote a custom Python implementation for PIC to keep track of the species contributing to each contrast, and to ensure transparency of method [71]. The ability to monitor the original species was useful when testing correlations on subsets of the entire phylogenetic tree.

### Analysis of vertebral data from extinct Theropods and Paraves

The vertebral counts for extinct Theropod and Paraves species were obtained from a number of literature sources which had analyzed the fossil remains [49, 50, 73–83]. We combined and averaged the counts for all taxa at the genus level. We used the phylogenetic tree or cladogram from Brusatte et al. [48] and manually trimmed it to the taxa for which we have the full vertebral count in the tree shown in Fig. 3c. We provide a table of all the vertebral data in the Supplementary Material as well as the corresponding source.

### BLASTing unannotated genomes

Many of the chromosome-level genome assemblies in the NCBI database have not yet been fully annotated. We determined the approximate location of the *Hox* genes by performing a BLAST search (blastn) [64] on each genome using the gene sequence from the corresponding *Hox* gene from a model organism in that tetrapod class as the search sequence. Thus for all unannotated bird genomes, for example, we used the chicken *Hox* genes to search for the genomic coordinates of their *Hox* genes. We then manually curated the search results and only retained species for which we were able to find all four *Hox* clusters with the entire cluster located on a single chromosome or scaffold. We include the list of annotated and unannotated genomes used in our study in the Supplementary Material, as well as the parameters used to perform the BLAST search. For the tuatara (*Sphenodon punctatus*) the top BLAST hit for *HoxB13* was significantly farther away from *HoxB9* then for other Squamata genomes so for this species and gene alone we searched using an alternative alignment tool, *progressiveMauve* [84].

### Determining the CTCF peak numbers

We tested the relationship between the number of CTCF binding sites and intergenic distances in the *Hox* clusters for several model organisms in different tetrapod classes. We used experimental ChIP-seq data from human H1 cells [85], mouse embryonic stem cells [86], chicken embryonic cells [87], and frog embryonic cells [88]. The human, mouse, and frog data were already analyzed and mapped to recent genome assemblies (GRCh38, GRCm38, and xenTro10). We mapped the raw reads for the chicken data to the bGalGal1 genome assembly using bowtie2 [89] with standard parameters and used samtools [90] with standard parameters to remove duplicates and sort the alignments. After smoothing the data we used the *find peaks* function from the Python *scipy* signal library [91] to detect peaks with a prominence larger than one standard deviation of the signal. Using the genome annotations from NCBI we determined the locations of the *Hox* genes in each cluster and counted the number of peaks in between each gene as well as determined the intergenic distance. We plot the sum of these counts and intergenic distances for each cluster and species in Extended Data Fig. 7a. The intergenic distances and number of CTCF binding sites have a Pearson correlation coefficient of *r* ≃ 0.81 and p-value ≃ 1.53 × 10^−4^.

### Silhouette and images

The silhouette images used in the figures were all obtained from www.phylopic.org except for *Hypotaenidia okinawae* which we created ourselves from an image obtained from the Ryuku Shimpo. The *Jeholornis* (*shenzhouraptor sinensis*) and *Hesperornis* (*regalis*) were created by Scott Hartman while the *Epidexipteryx* (for which we used *Scansoriopteryx heilmannii*) and *Sapeornis* (*chaoyangensis*) were created by Matthew Martyniuk under a Creative Commons license (https://creativecommons.org/ licenses/by/3.0/). All other images are in the public domain unless otherwise noted.

**Extended Data Figure 1.**
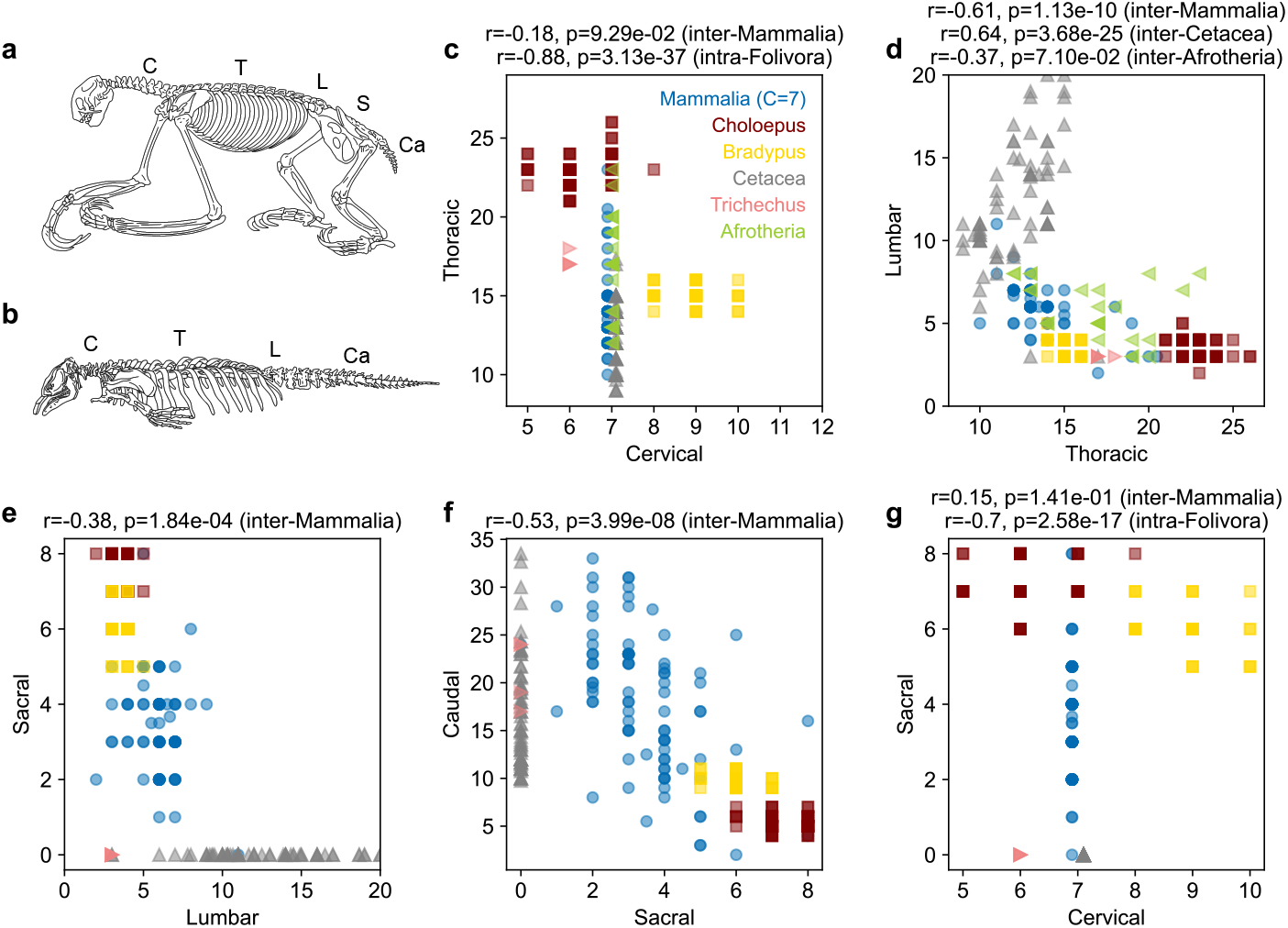
(a) Sketch of a three-toed sloth (*Bradypus tridactylus*) skeleton reproduced from [21]. (b) Sketch of a manatee (*Trichechus manatus*) skeleton reproduced from [92]. (c)-(f) Additional plots of adjacent vertebrae to explore homeotic patterns (category *II*) in Mammalia with additional data from many individual specimens of tree sloths (*Folivora*) [34], *Cetacea* [46], and *Afrotheria* (only *C, T*, and *L*) [23]. In (c) and (g) the *C* = 7 data points are slightly shifted for visibility. (c) A plot of *T* vs. *C* supports the hypothesis that within tree sloths (*Folivora*) these two vertebrae are related by a homeotic transformation as argued by [26]. Manatees also approximately follow the sloth trend. (d) Plot of *L* vs. *T* . Sloths and manatees follow the well-known mammalian constraint but Cetaceans exhibit an opposite trend, demonstrating that their *L* and *T* are not related by a homeotic transformation. Afrotherians show an increased amount of variation compared to other mammals [23]. We only plot to *L* = 20 to discern the variation of non-Cetaceans. (e)-(f) Plots of *S* vs. *L* and *Ca* vs. *S* shows that all mammals follow the newly discovered homeotic patterns. In (e) we only plot to *L* = 20 to discern the variation of non-Cetaceans. (g) Plot of *S* vs. *C*. Although an inter-species comparison within Mammalia only weakly follows the tetrapod distal balance constraint *C* − *S* ≃ 0, we find that an intra-*Folivora* comparison shows an opposite trend.

**Extended Data Figure 2.**
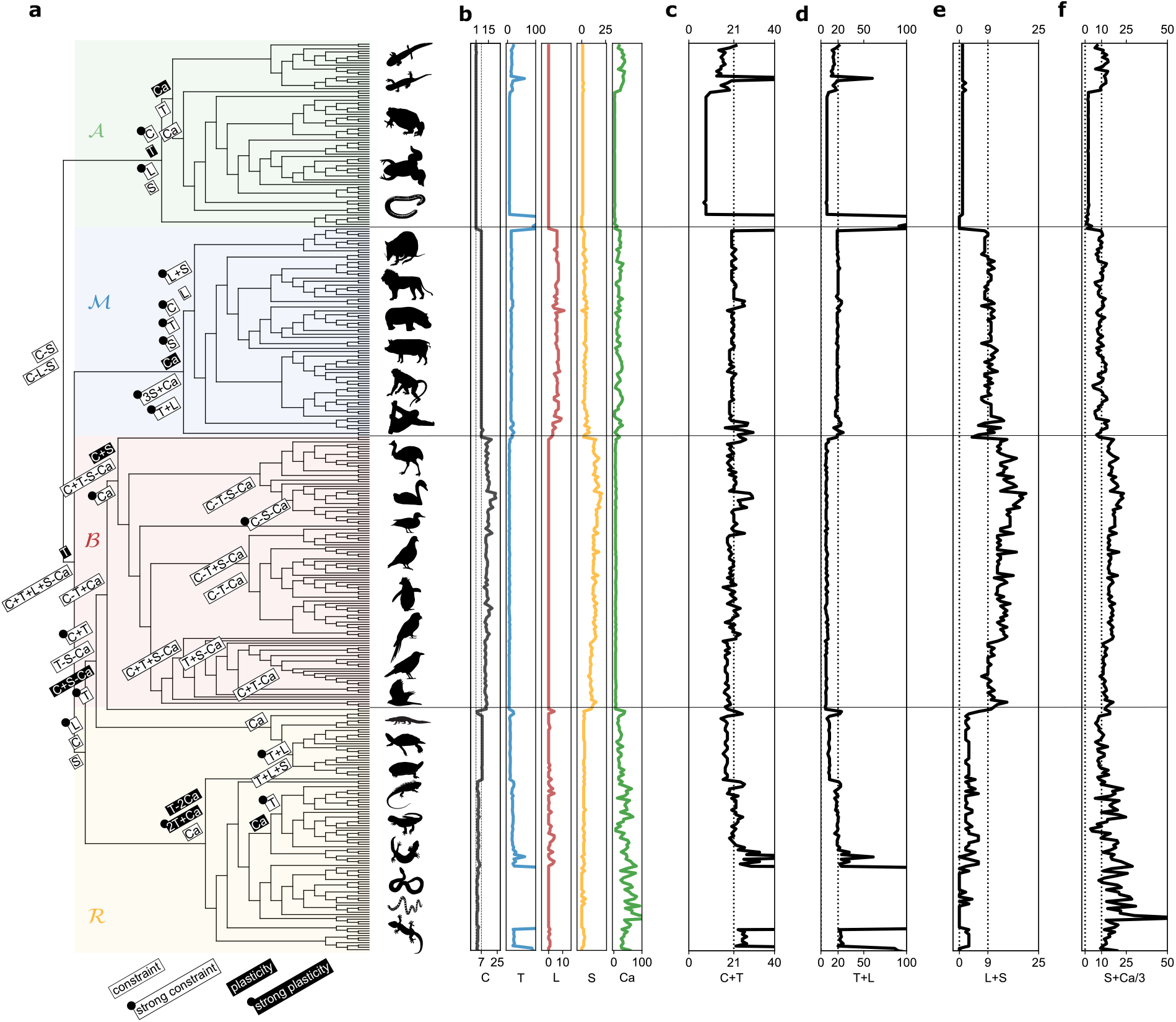
Extended plots of vertebral patterns using the phylogenetic tree to organize the vertical axis, with a focus on the homeotic patterns. (a) A phylogenetic tree containing all the species for which we have the full vertebral count. Unlike the tree in Fig. 1d, this version only shows topology. The tree has been organized so that the four classes of tetrapods, Amphibia, Mammalia, Aves (birds), and Reptiles, are arranged vertically. On top of the tree we label the branch *B* at which a constraint or plasticity has been identified where the rightmost point of the label corresponds to the branch point. We denote a “strong” constraint (plasticity) with a dot in the top left of the box. (b) The individual vertebrae plotted vertically. (c) Vertical plot of *C* + *T* . Excluding amphibians and snakes, most tetrapods follow a type-II constraint where *C* + *T* ≃ 21. (d) Vertical plot of *T* + *L*. Within mammals there is a type-II constraint where *T* + *L* ≃ 20. (Only mammals and several reptiles have *L* > 0.) (e) Vertical plot of *T* + *L*. Within mammals there is a type-II constraint where *L* + *S* ≃ 9. (Only mammals and several reptiles have *L* > 0.) (f) Vertical plot of *S* + *Ca*/3. Within mammals there is a type-II constraint where *S* + *Ca*/3 ≃ 10.

**Extended Data Figure 3.**
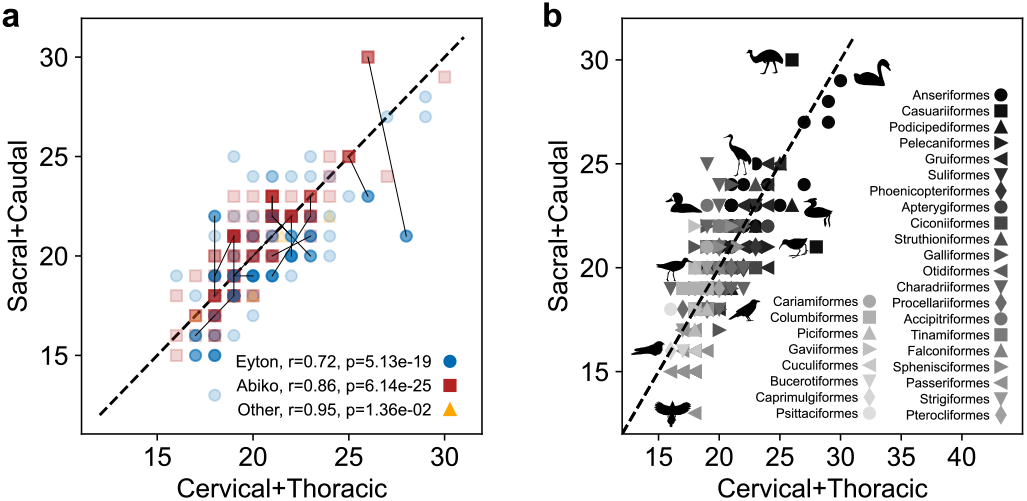
(a) A plot of *S* + *Ca* vs. *C* + *T* for all Aves specimens with the data source specified. Most data are from Eyton [22], another substantial amount is from Abiko Bird Museum, while the remainder are from various sources. When data from a single species comes from both Eyton and Abiko, we connected these data points by a solid line. Regardless of the data source, the adherence to the bird constraint remains essentially the same. (b) A plot of the posterior (*S* + *Ca*) of the bird vertebral counts vs. the anterior (*C* + *T* ) with the corresponding bird order represented by different markers. There are 29 unique orders. The shading of the markers is proportional to *C*. Within our dataset we find no significant order-specific deviation from the constraint.

**Extended Data Figure 4.**
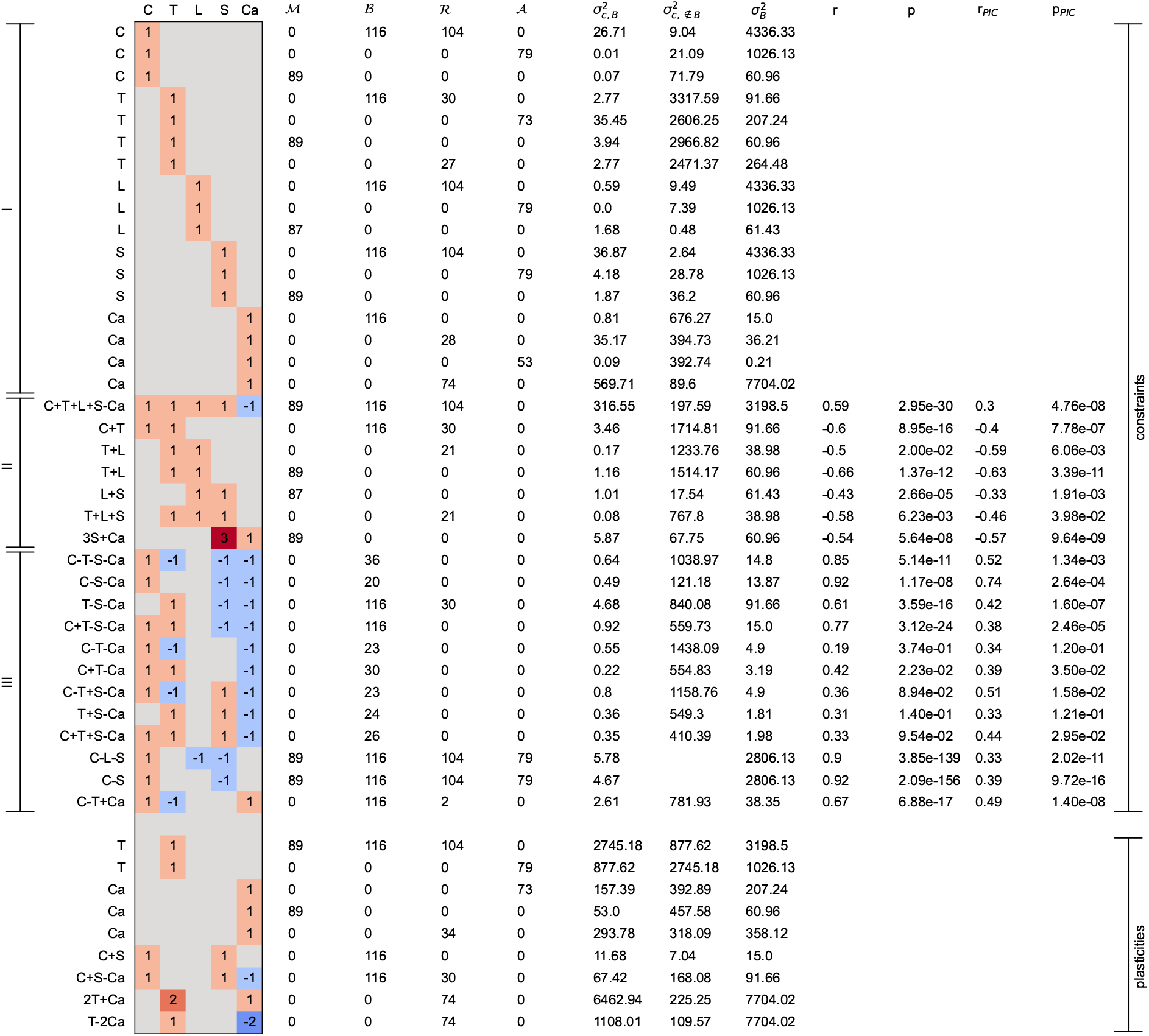
Table with all constraints and plasticities, including their formula, the number of species belonging to each tetrapod class, the associated variances, and the Pearson correlation coefficient (if calculable). We also denote the category of constraint (*I, II*, and *III*) as defined in the main text.

**Extended Data Figure 5.**
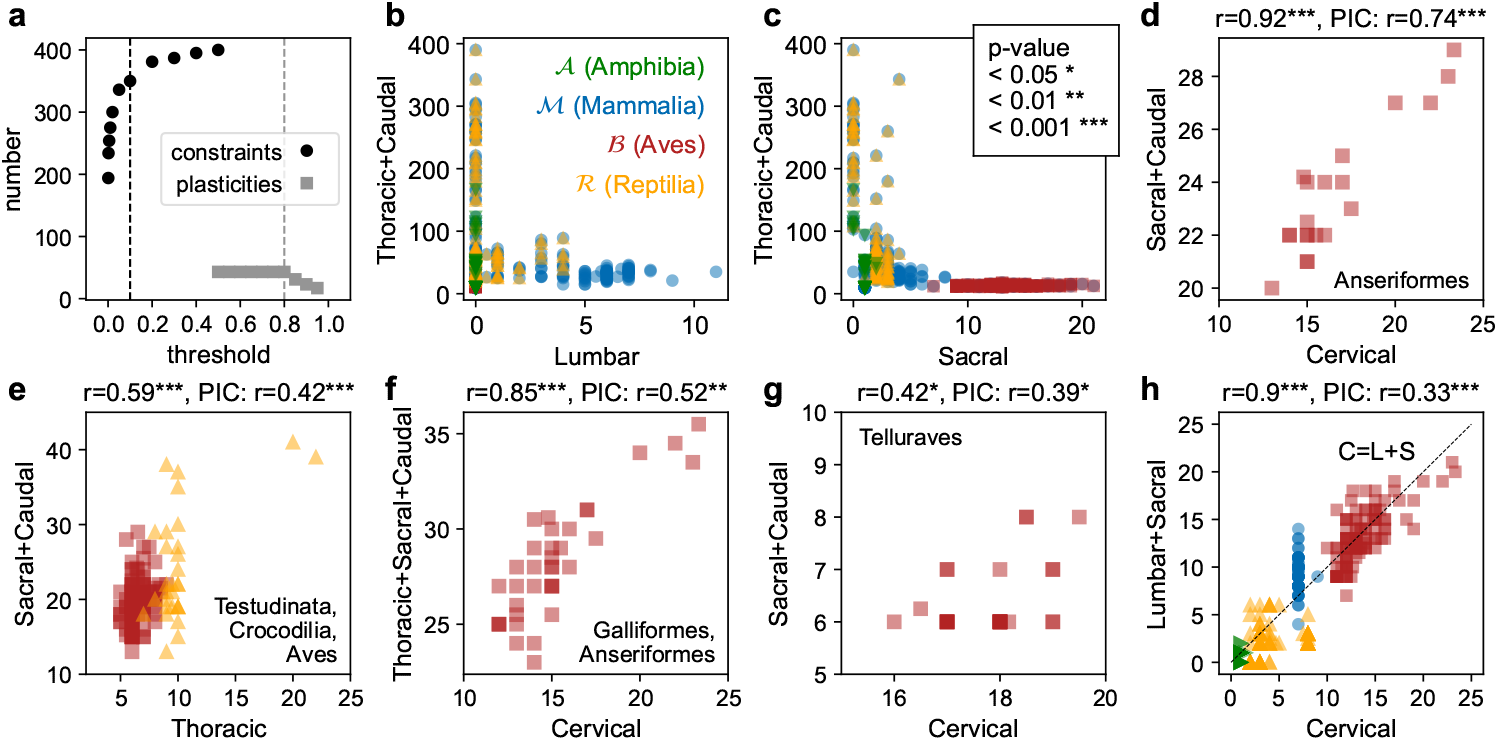
Plots of additional constraints, plasticities, and sensitivity to threshold. (a) Total number of constraints and plasticities (before combining and testing with PIC) vs. the threshold. For the constraints this is for both the ratio of the constraint variance inside the branch 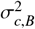 to the total variance of this branch 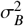 and with the variance of the same linear combination for all species (tips) outside this branch 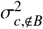 . For the plasticities it is just a threshold for the ratio 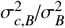 . The vertical dashed lines indicate the thresholds used to obtain the main results. In (b),(c) we provide additional examples of vertebrae which are constraints in one branch and plasticities in another, yielding a characteristic L-shape. All remaining constraints are a distal balance (category *III*). Many of the remaining constraints were found in sub-branches of Aves or with all of Aves and a sub-branch of Reptilia, and are slight variations on anterior-posterior bird constraint ((*C* + *T* ) − (*S* + *Ca*) ≃ 0). The distal balance shown in (h) is a slight variation of the *C* − *S* ≃ 0 constraint with the addition of *L* and is also at the tetrapod scale. For each plot where possible we note above it the Pearson correlation coefficient *r* before and after PIC [45] as well as the level of significance with *, **, or ***.

**Extended Data Figure 6.**
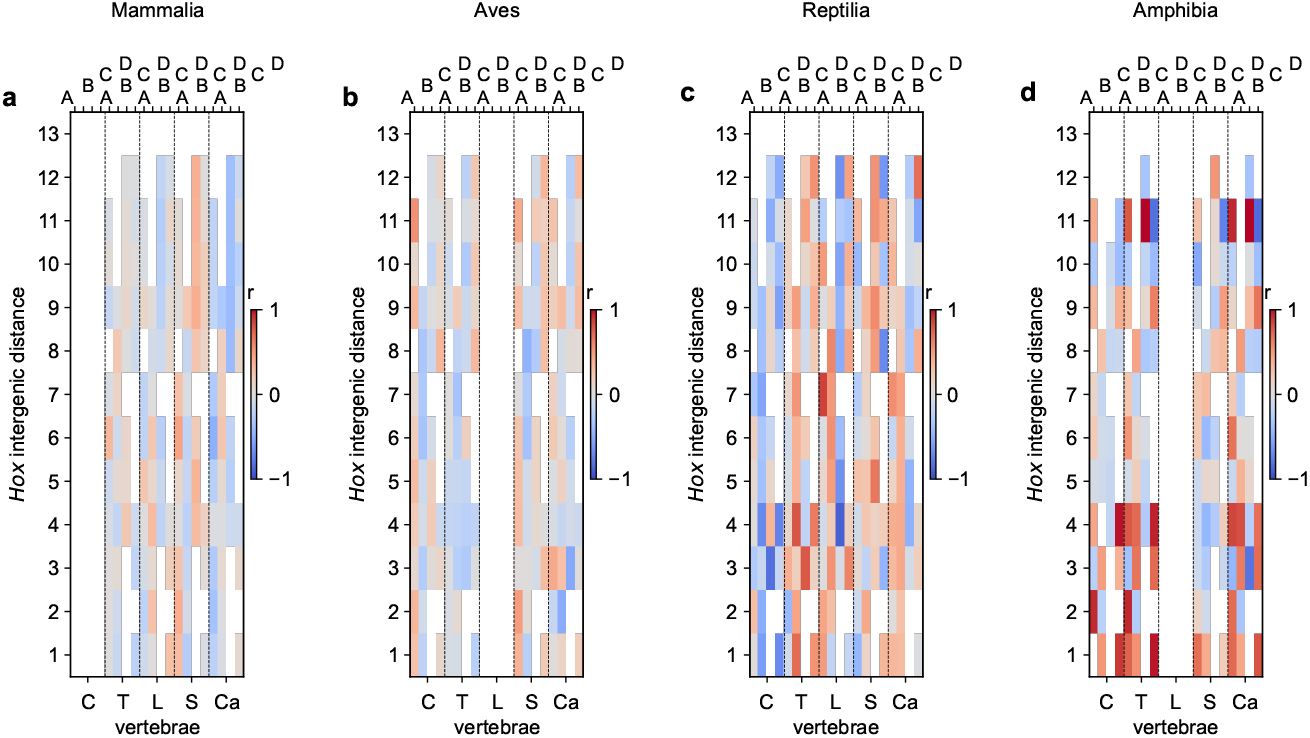
Plots of Pearson correlation coefficient matrix between *Hox* intergenic distances and vertebrae differentiated by tetrapod class. While there are individual *Hox* intergenic distances that correlate strongly with individual vertebal counts, an overall picture of colinearity does not emerge when examining individual classes.

**Extended Data Figure 7.**
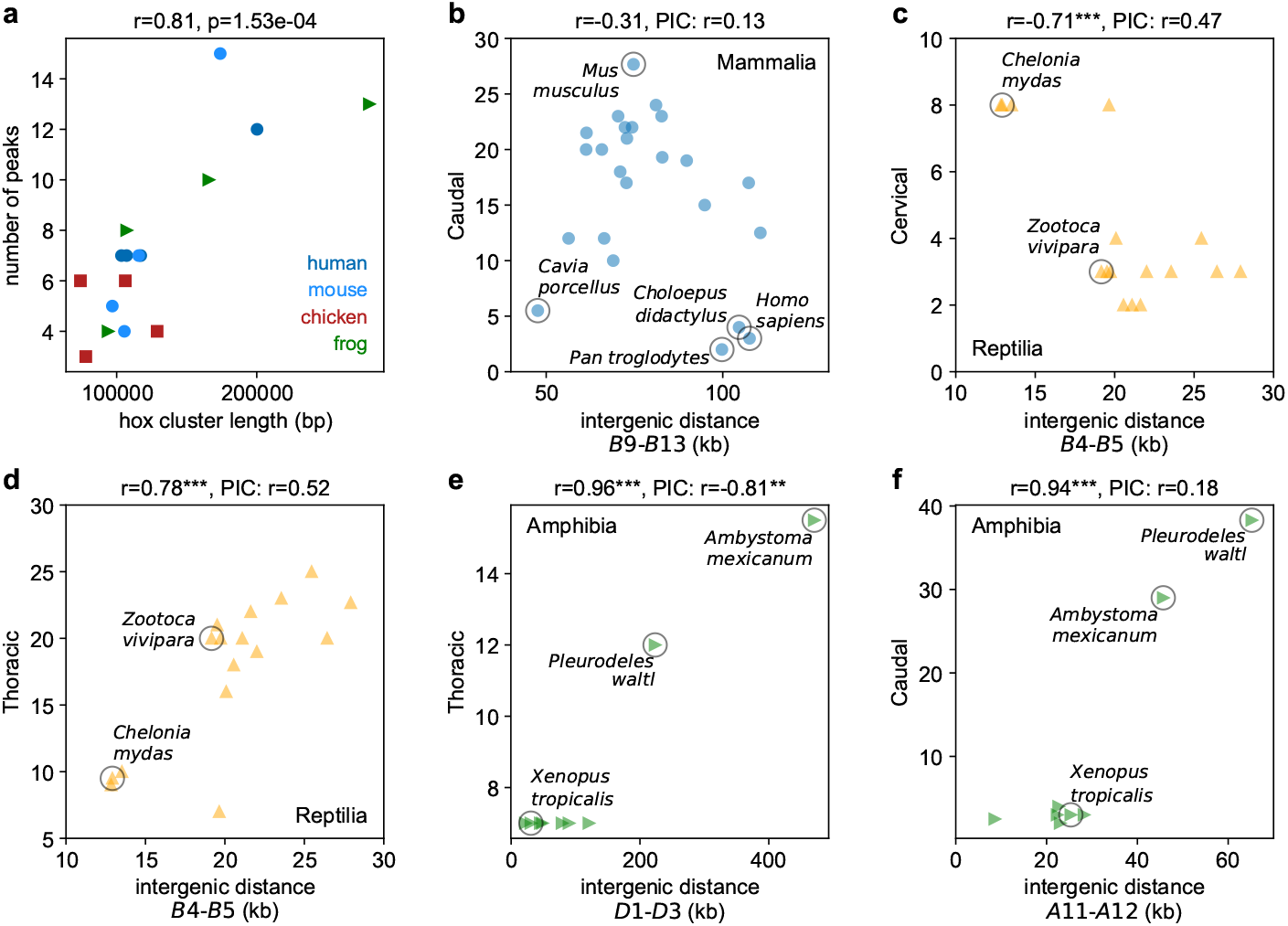
(a) Plot of experimental CTCF binding site counts vs. intergenic distance for different *Hox* clusters and species from several tetrapod classes. The Pearson correlation coefficient *r* is large and significant. (b)-(f) Additional plots of vertebral counts vs. intergenic distance with somewhat significant Pearson correlation coefficients *r*. While some individual *Hox* intergenic distances correlate strongly with individual vertebral counts, there does not seem to be strong evidence for an overall colinearity. Moreover, many of the stronger correlations do not survive the phylogenetic test (the correlation coefficient after PIC is small).

**Extended Data Figure 8.**
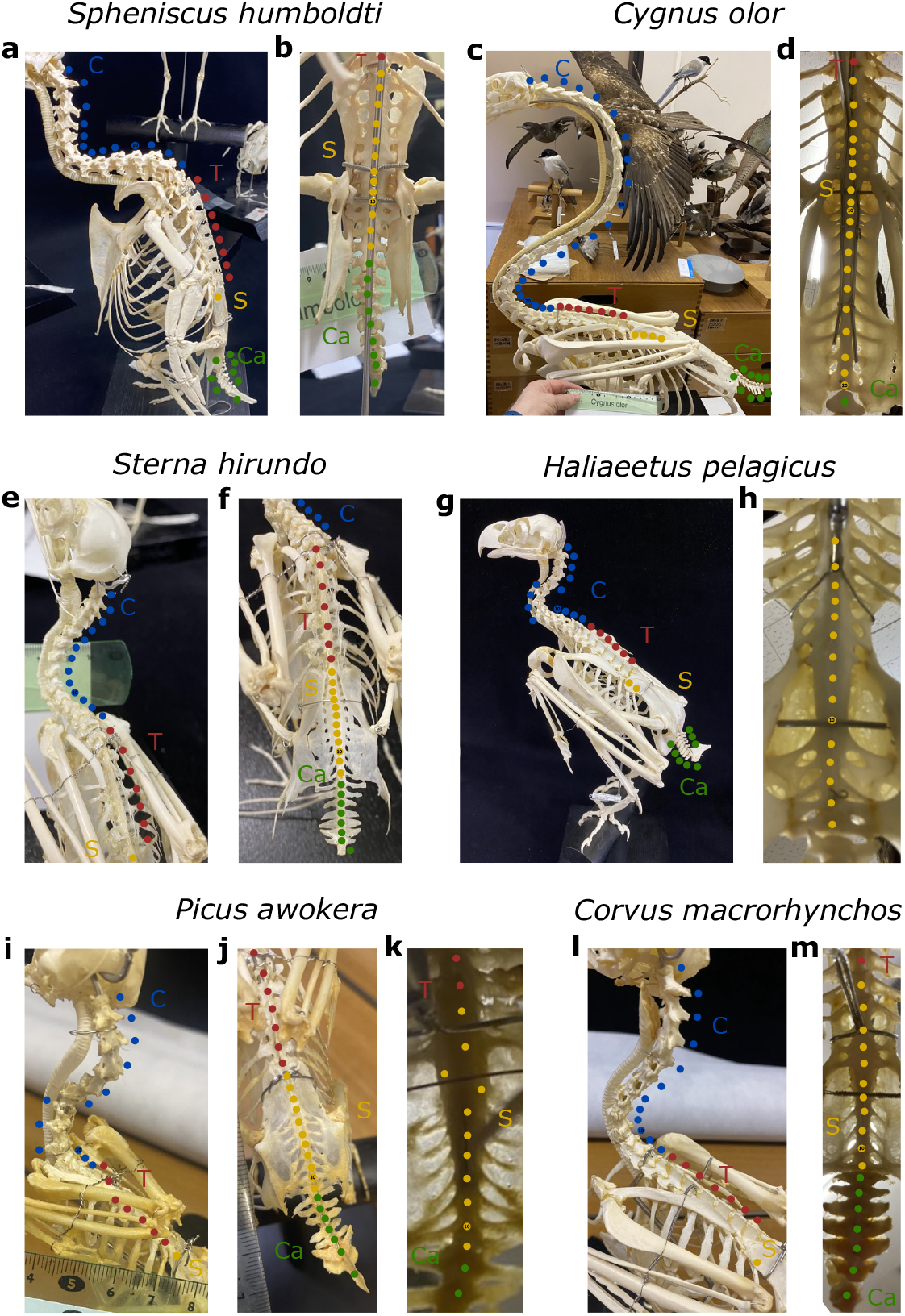
Several images of bird (Aves) skeletons from Abiko Bird Museum with the vertebrae indicated. In many cases, the individual vertebrae are difficult to distinguish in the central portion of the synsacrum. We relied on the presence of transverse processes (associated with canals whose proposed function is balance [93, 94]) and the size of preceding and following sacral vertebrae. (a)-(b) Photos of the skeleton of *Spheniscus humboldti* (Humboldt’s penguin). The sacral vertebrae are most often more easily discernible from an underside view as in (b). We determine the vertebral formula to be (13, 8, 0, 13, 9). (c)-(d) Photos of the skeleton of *Cygnus olor* (Mute swan). We determine the vertebral formula to be (23, 7, 0, 20, 9). (e)-(f) Photos of the skeleton of *Stern hirundo* (Common tern). Here the sacral vertebrae are distinguishable from an overhead view. We determine the vertebral formula to be (13, 7, 0, 12, 8). (g)-(h) Photos of the skeleton of *Haliaeetus pelagicus* (Steller’s sea eagle). We determine the vertebral formula to be (13, 6, 0, 15, 8). (i)-(k) Photos of the skeleton of *Picus awokera* (Japanese green woodpecker). We determine the vertebral formula to be (12, 7, 0, 11, 7). (l)-(m) Photos of the skeleton of *Corvus macrorhynchos* (Large-billed crow). We determine the vertebral formula to be (12, 7, 0, 11, 7).

**Extended Data Figure 9.**
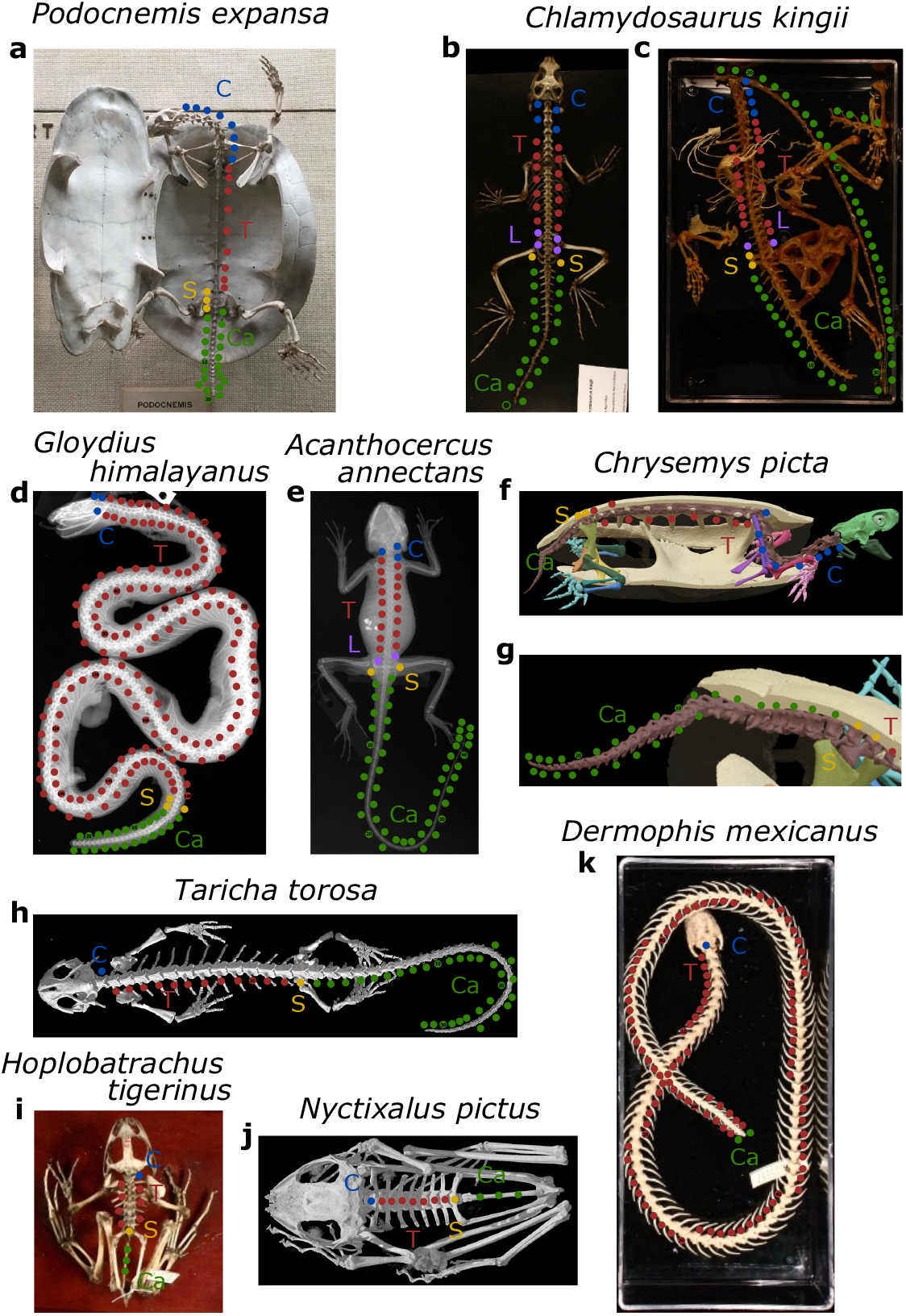
Several images of reptiles (Reptilia) and Amphibian skeletons from various sources with the vertebrae indicated. (a) Photo of the skeleton of *Podocnemis expansa* (Arrau turtle) retrieved from *iDigBio* and made available by Vertebrate Zoology Division of Herpetology at Yale Peabody Museum (YPM). We determine the vertebral formula to be (8, 9, 0, 3, 20). (b)-(c) Photos of the skeletons of two specimens of *Chlamydosaurus kingii* (Frilled lizard) retrieved from *iDigBio* and made available by YPM. The caudal series appears to be incomplete in (b) and the final few caudal vertebrae in (c) are difficult to distinguish so we use the lengths of the preceding vertebrae to estimate. Here we determine the consensus or average vertebral formula to be (4, 15, 3, 2, 52). (d) X-ray image of *Gloydius himalayanus* (Himalayan pit viper) retrieved from *iDigBio* and made available by the National Museum of Natural History, Smithsonian Institution (NMNH). The cloacal vertebrae (sacral) are identified by a shift in morphology from the preceding thoracic such as a reduction in the size or shape of the lymphapophyses (transverse processes) [65]. When it was not possible to identify these we set *S* = 0. Here we determine the vertebral formula to be (3, 158, 0, 3, 23). (e) X-ray image of *Acanthocercus annectans* (Eritrean rock agama) retrieved from *iDigBio* and made available by NMNH. Here we determine the vertebral formula to be (3, 17, 2, 2, 47). (f)-(g) Image of the CT scan of *Chrysemys picta* (Painted turtle) retrieved from *Sketchfab* and made available by Division of Amphibians and Reptiles, Museum of Southwestern Biology, University of New Mexico (MSB). Here we determine the vertebral formula to be (8, 10, 0, 2, 28). (h) Image of the CT scan of *Taricha torosa* (California newt) retrieved from *Digimorph* and made available by the Texas Science & Natural History Museum (TSNHM). We determine the vertebral formula to be (1, 12, 0, 1, 32). For *Anura* (frogs and toads), we follow the work of Ref. [72] who found that the Anuran tail (urostyle) likely comes from three to four vestigial caudal vertebrae as indicated by the presence of neural arches during embryonic development. We set the Anuran caudal count to be a uniform value of three. (i) Photo of the skeleton of *Hoplobatrachus tigerinus* (Indus valley bullfrog) retrieved from *iDigBio* and made available by YPM. We determine the vertebral formula to be (1, 7, 0, 1, 3). (j) Image of the CT scan of *Nyctixalus pictus* (Painted Indonesian treefrog) retrieved from *Digimorph* and made available by TSNHM. We determine the vertebral formula to be (1, 7, 0, 1, 3). (k) Photo of the skeleton of *Dermophis mexicanus* (Mexican burrowing caecilian) retrieved from *iDigBio* and made available by YPM. We determine the vertebral formula to be (1, 109, 0, 0, 2).

## References

[1] Müller, J. et al. Homeotic effects, somitogenesis and the evolution of vertebral numbers in recent and fossil amniotes. Proceedings of the National Academy of Sciences 107, 2118–2123 (2010).

[2] Mallo, M., Wellik, D. M. & Deschamps, J. Hox genes and regional patterning of the vertebrate body plan. Developmental biology 344, 7–15 (2010).

[3] Burke, A. C., Nelson, C. E., Morgan, B. A. & Tabin, C. Hox genes and the evolution of vertebrate axial morphology. Development 121, 333–346 (1995).

[4] Böhmer, C., Rauhut, O. W. & Wörheide, G. Correlation between hox code and vertebral morphology in archosaurs. Proceedings of the Royal Society B: Biological Sciences 282, 20150077 (2015).

[5] Woltering, J. M. et al. Axial patterning in snakes and caecilians: evidence for an alternative interpretation of the hox code. Developmental biology 332, 82–89 (2009).

[6] Woltering, J. M. & Duboule, D. Tetrapod axial evolution and developmental constraints; empirical underpinning by a mouse model. Mechanisms of development 138, 64–72 (2015).

[7] Ohya, Y. K., Kuraku, S. & Kuratani, S. Hox code in embryos of chinese soft-shelled turtle pelodiscus sinensis correlates with the evolutionary innovation in the turtle. Journal of Experimental Zoology Part B: Molecular and Developmental Evolution 304, 107–118 (2005).

[8] Böhmer, C. Correlation between hox code and vertebral morphology in the mouse: towards a universal model for synapsida. Zoological Letters 3, 1–11 (2017).

[9] Rashid, D. J. et al. From dinosaurs to birds: a tail of evolution. EvoDevo 5, 1–20 (2014).

[10] Di-Poi, N. et al. Changes in hox genes’ structure and function during the evolution of the squamate body plan. Nature 464, 99–103 (2010).

[11] Moreau, C. et al. Timed collinear activation of hox genes during gastrulation controls the avian forelimb position. Current Biology 29, 35–50 (2019).

[12] Narita, Y. & Kuratani, S. Evolution of the vertebral formulae in mammals: a perspective on developmental constraints. Journal of Experimental Zoology Part B: Molecular and Developmental Evolution 304, 91–106 (2005).

[13] Cuvier, G. Recherches sur les ossemens fossiles Vol. 5 (Dufour-D’Ocagne, 1825).

[14] Kumar, S. et al. Timetree 5: an expanded resource for species divergence times. Molecular Biology and Evolution 39, msac174 (2022).

[15] Belon, P. & Glardon, P. L’histoire de la nature des oyseaux 306 (Librairie Droz, 1997).

[16] Cole, F. J. & Eales, N. B. The history of comparative anatomy: Part i.—a statistical analysis of the literature. Science Progress (1916-1919) 11, 578–596 (1917).

[17] Saint-Hilaire, É. G. Cours de l’histoire naturelle des Mamifères (Pichon, 1829).

[18] baron Cuvier, G. Leçons d’anatomie compareé Vol. 3 (Baudouin, 1840).

[19] Owen, R. Descriptive catalogue of the osteological series contained in the museum Vol. 2 (Taylor & Francis, 1853).

[20] Owen, R. Anatomy of Vertebrates, Vol. 1: Fishes and Reptiles (Longmans, Green, and Company, London, 1866).

[21] Owen, R. On the anatomy of vertebrates: birds and mammals Vol. 2 (Longmans, Green and Company, 1866).

[22] Eyton, T. C. Osteologia Avium: Or, A Sketch of the Osteology of Birds Vol. 1 (R. Hobson, 1875).

[23] Sánchez-Villagra, M. R., Narita, Y. & Kuratani, S. Thoracolumbar vertebral number: the first skeletal synapomorphy for afrotherian mammals. Systematics and Biodiversity 5, 1–7 (2007).

[24] Galis, F. et al. Fast running restricts evolutionary change of the vertebral column in mammals. Proceedings of the National Academy of Sciences 111, 11401–11406 (2014).

[25] Williams, S. A. et al. Increased variation in numbers of presacral vertebrae in suspensory mammals. Nature ecology & evolution 3, 949–956 (2019).

[26] Varela-Lasheras, I. et al. Breaking evolutionary and pleiotropic constraints in mammals: on sloths, manatees and homeotic mutations. EvoDevo 2, 1–27 (2011).

[27] He, S. et al. An axial hox code controls tissue segmentation and body patterning in nematostella vectensis. Science 361, 1377–1380 (2018).

[28] Liang, D., Wu, R., Geng, J., Wang, C. & Zhang, P. A general scenario of hoxgene inventory variation among major sarcopterygian lineages. BMC evolutionary biology 11, 1–13 (2011).

[29] Duboule, D. The (unusual) heuristic value of hox gene clusters; a matter of time? Developmental Biology (2022).

[30] Kessel, M. & Gruss, P. Homeotic transformations of murine vertebrae and concomitant alteration of hox codes induced by retinoic acid. Cell 67, 89–104 (1991).

[31] Alberch, P. From genes to phenotype: dynamical systems and evolvability. Genetica 84, 5–11 (1991).

[32] Pigliucci, M. Genotype–phenotype mapping and the end of the ‘genes as blueprint’metaphor. Philosophical Transactions of the Royal Society B: Biological Sciences 365, 557–566 (2010).

[33] Duboule, D. & Morata, G. Colinearity and functional hierarchy among genes of the homeotic complexes. Trends in Genetics 10, 358–364 (1994).

[34] Buchholtz, E. A. & Stepien, C. C. Anatomical transformation in mammals: developmental origin of aberrant cervical anatomy in tree sloths. Evolution & development 11, 69–79 (2009).

[35] Hautier, L., Weisbecker, V., Sánchez-Villagra, M. R., Goswami, A. & Asher, R. J. Skeletal development in sloths and the evolution of mammalian vertebral patterning. Proceedings of the National Academy of Sciences 107, 18903–18908 (2010).

[36] Böhmer, C., Amson, E., Arnold, P., van Heteren, A. H. & Nyakatura, J. A. Homeotic transformations reflect departure from the mammalian ‘rule of seven’cervical vertebrae in sloths: inferences on the hox code and morphological modularity of the mammalian neck. BMC Evolutionary Biology 18, 1–11 (2018).

[37] Szczygielski, T. Homeotic shift at the dawn of the turtle evolution. Royal Society Open Science 4, 160933 (2017).

[38] Mao, F., Zhang, C., Liu, C. & Meng, J. Fossoriality and evolutionary development in two cretaceous mammaliamorphs. Nature 592, 577–582 (2021).

[39] Figueirido, B. et al. Body-axis organization in tetrapods: a model-system to disentangle the developmental origins of convergent evolution in deep time. Biology Letters 18, 20220047 (2022).

[40] Li, Y. et al. Divergent vertebral formulae shape the evolution of axial complexity in mammals. Nature Ecology & Evolution 7, 367–381 (2023).

[41] Böhmer, C., Plateau, O., Cornette, R. & Abourachid, A. Correlated evolution of neck length and leg length in birds. Royal Society open science 6, 181588 (2019).

[42] Smith, J. M. et al. Developmental constraints and evolution: a perspective from the mountain lake conference on development and evolution. The Quarterly Review of Biology 60, 265–287 (1985).

[43] Antonovics, J. & van Tienderen, P. H. Ontoecogenophyloconstraints? the chaos of constraint terminology. Trends in Ecology & Evolution 6, 166–168 (1991).

[44] Owen, R. On the nature of limbs: a discourse delivered on Friday, February 9, at an Evening Meeting of the Royal Institution of Great Britain (J. van Voorst, 1849).

[45] Felsenstein, J. Phylogenies and the comparative method. The American Naturalist 125, 1–15 (1985).

[46] Buchholtz, E. A. & Gee, J. K. Finding sacral: developmental evolution of the axial skeleton of odontocetes (cetacea). Evolution & Development 19, 190–204 (2017).

[47] Ringnér, M. What is principal component analysis? Nature biotechnology 26, 303–304 (2008).

[48] Brusatte, S. L., Lloyd, G. T., Wang, S. C. & Norell, M. A. Gradual assembly of avian body plan culminated in rapid rates of evolution across the dinosaur-bird transition. Current Biology 24, 2386–2392 (2014).

[49] Zhang, F., Zhou, Z., Xu, X., Wang, X. & Sullivan, C. A bizarre jurassic maniraptoran from china with elongate ribbon-like feathers. Nature 455, 1105–1108 (2008).

[50] Clarke, J. A., Zhou, Z. & Zhang, F. Insight into the evolution of avian flight from a new clade of early cretaceous ornithurines from china and the morphology of yixianornis grabaui. Journal of anatomy 208, 287–308 (2006).

[51] Gao, C. et al. A new basal lineage of early cretaceous birds from china and its implications on the evolution of the avian tail. Palaeontology 51, 775–791 (2008).

[52] Brusatte, S. L., O’Connor, J. K. & Jarvis, E. D. The origin and diversification of birds. Current Biology 25, R888–R898 (2015).

[53] Wang, X. et al. New evidence from china for the nature of the pterosaur evolutionary transition. Scientific Reports 7, 42763 (2017).

[54] O’Connor, J. Enantiornithes. Current Biology 32, R1166–R1172 (2022).

[55] Walton, D. W. & Walton, G. M. Post-cranial osteology of bats. Fondren Science Series 1, 7 (1970).

[56] Igado, O. O. & Ade-Julius, E. R. Gross description and osteometrics of the axial skeleton (ribs and vertebrae) of eidolon helvum (african fruit bat). Nigerian Journal of Physiological Sciences 33, 189–194 (2018).

[57] Williston, S. W. On the osteology of nyctosaurus (nyctodactylus), with notes on american pterosaurs. Field Columbian Museum Geological Series 2 (1903).

[58] Bennett, S. C. The osteology and functional morphology of the late cretaceous pterosaur pteranodon part i. general description of osteology. Palaeontographica Abteilung A 1–112 (2001).

[59] Unwin, D. M., Lü, J. & Bakhurina, N. N. On the systematic and stratigraphic significance of pterosaurs from the lower cretaceous yixian formation (jehol group) of liaoning, china. Fossil Record 3, 181–206 (2000).

[60] Hone, D., Henderson, D. M., Therrien, F. & Habib, M. B. A specimen of rhamphorhynchus with soft tissue preservation, stomach contents and a putative coprolite. PeerJ 3, e1191 (2015).

[61] Ősi, A. & Prondvai, E. Forgotten pterosaurs in hungarian collections: first description of rhamphorhynchus and pterodactylus specimens. Neues Jahrbuch für Geologie und Palaöntologie-Abhandlungen 167–180 (2009).

[62] Rekaik, H. et al. Sequential and directional insulation by conserved ctcf sites underlies the hox timer in stembryos. Nature Genetics 1–12 (2023).

[63] Deschamps, J. & Duboule, D. Embryonic timing, axial stem cells, chromatin dynamics, and the hox clock. Genes & development 31, 1406–1416 (2017).

[64] Altschul, S. F., Gish, W., Miller, W., Myers, E. W. & Lipman, D. J. Basic local alignment search tool. Journal of molecular biology 215, 403–410 (1990).

[65] Gomez, C. et al. Control of segment number in vertebrate embryos. Nature 454, 335–339 (2008).

[66] Yamanaka, Y. et al. Reconstituting human somitogenesis in vitro. Nature 614, 509–520 (2023).

[67] Galis, F. Why do almost all mammals have seven cervical vertebrae? developmental constraints, hox genes, and cancer. Journal of Experimental Zoology 285, 19–26 (1999).

[68] Buchholtz, E.A. et al. Fixed cervical count and the origin of the mammalian diaphragm. Evolution & development 14, 399–411 (2012).

[69] Bui, H.-N. N.& Larsson, H. C. Development and evolution of regionalization within the avian axial column. Zoological Journal of the Linnean Society 191, 302–321 (2021).

[70] Lindsey, C. Pleomerism, the widespread tendency among related fish species for vertebral number to be correlated with maximum body length. Journal of the Fisheries Board of Canada 32, 2453–2469 (1975).

[71] Repository for supplementary data and source code.

[72] Ročková, H. & Roček, Z. Development of the pelvis and posterior part of the vertebral column in the anura. Journal of Anatomy 206, 17–35 (2005).

[73] Ostrom, J. H. Archaeopteryx and the origin of birds. Biological Journal of the linnean Society 8, 91–182 (1976).

[74] Chiappe, L. M., Ji, S.-A., Ji, Q. & Norell, M. A. Anatomy and systematics of the confuciusornithidae (theropoda, aves) from the late mesozoic of northeastern china. bulletin of the amnh; no. 242 (1999).

[75] Zhou, Z. & Zhang, F. Jeholornis compared to archaeopteryx, with a new understanding of the earliest avian evolution. Naturwissenschaften 90, 220–225 (2003).

[76] Pei, R., Li, Q., Meng, Q., Norell, M. A. & Gao, K.-Q. New specimens of anchiornis huxleyi (theropoda: Paraves) from the late jurassic of northeastern china. Bulletin of the American Museum of Natural History 2017, 1–67 (2017).

[77] Ostrom, J. H. & Gauthier, J. A. Osteology of Deinonychus antirrhopus, an unusual theropod from the Lower Cretaceous of Montana (Yale University Press, 2019).

[78] Zhong-He, Z., Xiao-Lin, W., Fu-Cheng, Z. & Xing, X. Important features of caudipteryxevidence from two nearly complete new specimens. Vertebrata PalAsiatica 38, 241 (2000).

[79] Qiang, J., Currie, P. J., Norell, M. A. & Shu-An, J. Two feathered dinosaurs from northeastern china. Nature 393, 753–761 (1998).

[80] Brochu, C. A. Osteology of tyrannosaurus rex: insights from a nearly complete skeleton and high-resolution computed tomographic analysis of the skull. Journal of vertebrate Paleontology 22, 1–138 (2003).

[81] Benito, J. et al. Forty new specimens of ichthyornis provide unprecedented insight into the postcranial morphology of crownward stem group birds. PeerJ 10, e13919 (2022).

[82] Bell, A. & Chiappe, L. M. Anatomy of parahesperornis: evolutionary mosaicism in the cretaceous hesperornithiformes (aves). Life 10, 62 (2020).

[83] Norell, M. A. & Clarke, J. A. Fossil that fills a critical gap in avian evolution. Nature 409, 181–184 (2001).

[84] Darling, A. E., Mau, B. & Perna, N. T. pro-gressivemauve: multiple genome alignment with gene gain, loss and rearrangement. PloS one 5, e11147 (2010).

[85] Zhang, J. et al. An integrative encode resource for cancer genomics. Nature communications 11, 3696 (2020).

[86] Bonev, B. et al. Multiscale 3d genome rewiring during mouse neural development. Cell 171, 557–572 (2017).

[87] Kadota, M. et al. Ctcf binding landscape in jawless fish with reference to hox cluster evolution. Scientific reports 7, 1–11 (2017).

[88] Niu, L. et al. Three-dimensional folding dynamics of the xenopus tropicalis genome. Nature Genetics 53, 1075–1087 (2021).

[89] Langmead, B. & Salzberg, S. L. Fast gapped-read alignment with bowtie 2. Nature methods 9, 357–359 (2012).

[90] Li, H. et al. The sequence alignment/map format and samtools. bioinformatics 25, 2078–2079 (2009).

[91] Virtanen, P. et al. Scipy 1.0: fundamental algorithms for scientific computing in python. Nature methods 17, 261–272 (2020).

[92] Knight, C. The pictorial museum of animated nature (London: C. Cox, 1844).

[93] Necker, R. Specializations in the lumbosacral vertebral canal and spinal cord of birds: evidence of a function as a sense organ which is involved in the control of walking. Journal of Comparative Physiology A 192, 439–448 (2006).

[94] Stanchak, K. E., French, C., Perkel, D. J. & Brunton, B. W. The balance hypothesis for the avian lumbosacral organ and an exploration of its morphological variation. Integrative Organismal Biology 2, obaa024 (2020).

